# Structure and mechanism of the human TMEM260 O-mannosyltransferase

**DOI:** 10.64898/2026.03.17.711096

**Authors:** Javier O. Cifuente, Lorenzo Povolo, Borja Ochoa-Lizarralde, Serpil Ahmed, Sergey Y. Vakhrushev, Jorge P. López-Alonso, Igor Tascón, Javier Fernandez-Martinez, Hiren J. Joshi, Adnan Halim, Iban Ubarretxena-Belandia

## Abstract

Protein O-linked mannose (O-Man) glycosylation is essential for mammalian development, and mutations in its biosynthetic glycosyltransferases cause severe muscular, neurological and cardiac disorders. Despite its biological and clinical importance, the structural basis of mammalian O-Man biosynthesis has remained unknown. Here we report cryo-electron microscopy (cryo-EM) structures of human TMEM260, an endoplasmic reticulum glycosyltransferase that selectively catalyzes O-mannosylation of semaphorin plexin receptors and receptor tyrosine kinases cMET and RON, key regulators of cell guidance and migration. Structures of TMEM260 in a ternary complex with its natural donor dolichyl-phosphate-β-mannose (Dol-P-Man) and an acceptor peptide derived from plexin-B2, together with binary complexes with Dol-P-Man or a synthetic donor analogue, capture physiologically relevant, substrate-loaded states and reveal the structural basis of O-Man transfer. We identify a conserved O-mannosylation sequon that underlies acceptor specificity and show that TMEM260 modifies extended polypeptide substrates, consistent with a co-translational glycosylation mechanism. These findings establish the molecular mechanism of a mammalian O-mannosyltransferase required for the maturation of physiologically critical receptors and provide a structural framework for interpreting TMEM260-associated congenital malformations.

## Introduction

Protein glycosylation covalently appends carbohydrate moieties (glycans) to proteins, affecting their folding, stability, trafficking and function [1]. Along the secretory pathway, specialized glycosyltransferases (GTs) catalyze site-specific N-, C- and O-linked glycan biosynthesis, generating structural and functional diversity essential for eukaryotic life [2–4]. Accordingly, defects in GTs underlie a growing spectrum of congenital disorders of glycosylation (CDG), which often manifest as severe multisystem developmental abnormalities [5, 6].

O-Man glycosylation initiated in the endoplasmic reticulum (ER) is among the most complex, physiologically important, and clinically relevant O-glycosylation pathways in mammals [5, 6]. However, the structural basis of mammalian O-Man biosynthesis has remained unknown. The molecular mechanism governing donor and acceptor recognition, O-Man transfer, and the overall architecture of enzymes initiating O-Man biosynthesis are unresolved. Three families of O-mannosyltransferases, POMT1/2, TMTC1–4 and TMEM260, initiate O-Man biosynthesis on distinct classes of substrate proteins [7]. The POMT1/2 pathway acting on alpha-dystroglycan (αDG) established the molecular basis of dystroglycanopathies caused by defective matriglycan synthesis [8, 9]. TMTC1–4 isoenzymes were subsequently shown to glycosylate cadherins and protocadherins, linking their dysfunction to neurodevelopmental defects, including hearing loss and severe brain malformations [10–12].

TMEM260 defines the most recently identified O-Man initiation pathway, transferring α-mannose from Dol-P-Man onto specific serine or threonine residues (**Fig. S1a**) within Immunoglobulin-like fold, Plexin, Transcription-factors (IPT) domains of the hepatocyte growth factor receptor (cMET), macrophage-stimulating protein receptor (RON), and semaphorin plexin family receptors involved in cell guidance and migration (**Fig. S1b; Table S1**) [13]. Genetic defects in *TMEM260* cause (**Table S2**) structural heart defects and renal anomalies (SHDRA; OMIM4 617478), an autosomal-recessive CDG characterized by congenital heart malformations, including persistent truncus arteriosus (PTA), and high infant mortality [13–21]. Recent genetic surveys suggest that TMEM260-related congenital heart disease may represent a major cause of PTA in Japan [19, 21]. At the cellular level, loss of TMEM260-dependent O-Man transfer impairs receptor maturation and signalling, leading to abnormal epithelial morphogenesis [13].

Here, we present cryo-electron microscopy (cryo-EM) structures of human TMEM260 in a ternary complex (3.1 Å resolution) with the natural donor Dol-P-Man and an acceptor peptide from the IPT1 domain of the plexin-B2 receptor (PLXNB2), and in binary complexes with Dol-P-Man (2.9 Å) and the synthetic donor analogue farnesyl-phosphate-β-mannose (Far-P-Man; 2.7 Å). The structures capture donor- and acceptor-loaded states, and define the mechanism of substrate recognition and O-Man transfer. Together, they establish a framework for understanding mammalian O-Man glycosylation and for interpreting TMEM260 variants that underlie congenital developmental disorders.

## Results

### A transmembrane GT-C core coupled to ER-luminal Rossmann-like and TPR domains

We set out to define the architecture of human TMEM260 (**Fig. S2a**) and how it binds glycolipid donor. Because the natural donor Dol-P-Man (**Fig. S2d**) is not commercially available, we incubated the purified enzyme (**Fig. S2b-c**) with an excess of the synthetic donor analogue Far-P-Man (**Fig. S2e; Table S3**). Single-particle cryo-EM reconstruction at a nominal resolution of 2.7 Å (**Fig. 1a; Table S4; Figs. S3a**,**S4a**) resolved residues 24–706 of TMEM260 in a DDM micelle bound to a Far-P-Man molecule (**Fig. 1b; PDB 9RQL; Fig. S5**), with density visible for the complete isoprenoid chain (**Fig. S6a**).

**Fig. 1.**
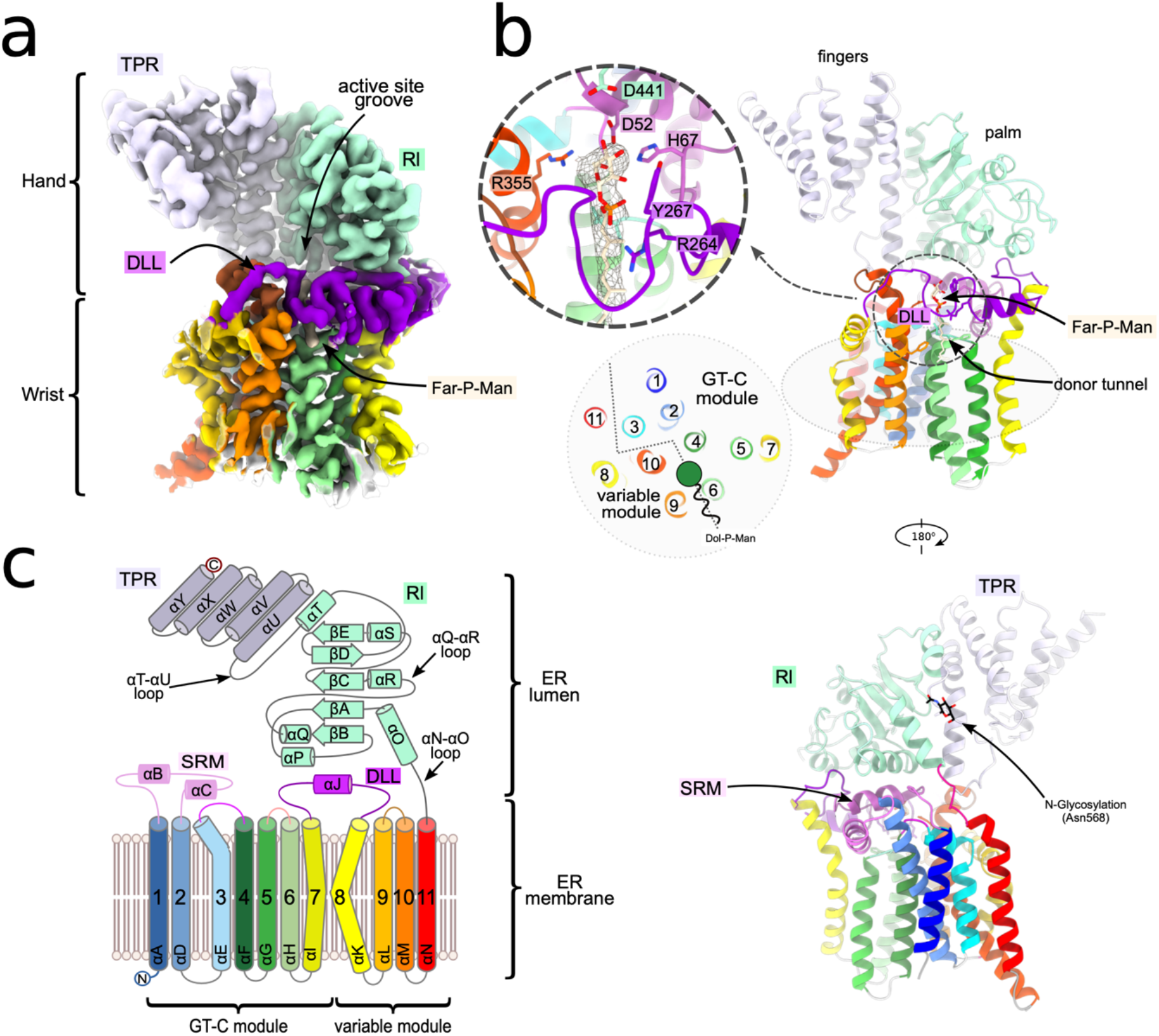
Structure of TMEM260 bond to the donor analogue Far-P-Man. **a**, Cryo-EM density map of TMEM260 in complex with the synthetic donor analogue Far-P-Man, determined at 2.7 Å resolution. Major structural regions are indicated and colored consistently throughout the figure. **b**, Front (top) and back (bottom) views of the corresponding atomic model shown in ribbon representation. Close-up (dashed circle) highlights the donor-analogue density and fitted model, with interacting residues annotated. A schematic section parallel to the membrane plane (grey plane) illustrates the arrangement of the transmembrane helices within the GT-C and variable modules. The position of Far-P-Man is depicted schematically, with the sugar moiety shown as a green circle and the phosphate and hydrophobic chain represented as a curved tail. **c**, Topology diagram depicting the arrangement of α-helices (cylinders), β-strands (arrows) and connecting loops, labelled from N to the C terminus. Transmembrane segments are numbered sequentially, and conserved motifs are indicated.

Based on their structural folds, GTs are classified into four major classes: GT-A, GT-B, GT-C, and lysozyme-like [22]. GT-A, GT-B, and lysozyme-like enzymes are soluble, membrane-associated, or bitopic membrane proteins. In contrast, members of the GT-C superfamily are multi-pass transmembrane proteins. Consistent with this classification, TMEM260 displays a GT-C fold composed of two transmembrane modules (residues 24–415) [23], a GT-C module formed by seven transmembrane helices (TMHs 1–7) and a variable module (VM) comprising four helices (TMHs 8–11), which connects to an ER-luminal lobe containing Rossmann-like (Rl, residues 416–564) and tetratricopeptide-repeat (TPR, residues 565–706) domains (**Fig. 1c**). The enzyme adopts an articulated molecular “hand” architecture (**Fig. 1a,b**) where the eleven TMHs form a compact donor-binding “wrist” anchoring TMEM260 in the ER membrane. The Rl domain forms the “palm” of the molecular hand and provides the central structural platform, while the distal TPR domain comprises finger-like helices projecting from the palm.

The GT-C module contains a sugar-recognition motif (SRM; residues 45–90) between TMHs 1 and 2, comprising helices αB-αC and loops that shape the active site and connect the wrist to the ER-luminal lobe. The GT-C module ends at TMH 7, which connects to TMH 8 of the VM via a membrane-parallel bridge helix (αJ). A donor-loading loop (DLL; residues 263–279) arches over TMHs 6 and 9 and delineates the donor tunnel. TMH 8 deviates from canonical α-helical geometry, splitting into an amphipathic segment and a transmembrane segment. TMHs 4 and 6 pack against TMHs 9 and 10, forming the transmembrane core that houses Far-P-Man, with luminal termini defining the active-site cavity at the membrane-lumen interface. TMH 11 adopts a quasi-perpendicular orientation, linking the transmembrane and luminal regions. The ER-luminal lobe comprises both Rl and TPR domains. The Rl domain forms the palm, built around a five-stranded central β-sheet flanked by α-helices organized into two subregions that rest above the GT-C wrist and shape a cavity aligned with the catalytic site. An interdomain linker (N568 is N-glycosylated, **Fig. 1b**) connects the palm to a canonical five-helix TPR fold that forms the fingers.

The Far-P-Man β-mannose ring lies in the energetically favored ^4^C_1_ chair conformation (**Fig. 1b; Fig. S6a**). Conserved SRM residues D52 and H67, Y267 in the DLL and R355 in TMH 10 engage the mannose, the phosphate group interacts with R264, and the farnesyl chain extends into the hydrophobic donor tunnel in the wrist. To visualize the apo state, we purified TMEM260 without exogenous substrates and determined its structure (**Figs. S3b**,**S4b; PDB 9RQN; Table S4**). Unexpectedly, the 2.9 Å map contained clear density for endogenous Dol-P-Man, resolved up to the fifteenth carbon of its isoprenoid chain (**Fig. S6b**), indicating co-purification from HEK293 membranes. Dol-P-Man occupies the same binding site as Far-P-Man, with the sugar in a ^4^C_1_ chair conformation and the dolichyl chain extending into the donor tunnel beneath the DLL.

DALI [24] searches of the isolated Rl and TPR domains retrieved similarities to known GTs (**Fig. S7**), consistent with evolutionary relationships at the domain level. By contrast, queries using the full-length structure did not retrieve hits among GTs or other membrane proteins, underscoring the uniqueness of the integrated molecular hand architecture of TMEM260. Across species, strictly conserved residues cluster predominantly within the transmembrane wrist and active-site core, with conservation progressively decreasing toward the distal TPR fingers (**Fig. S8**), highlighting the primacy of the transmembrane-luminal interface in TMEM260 function.

### O-mannosylation of PLXNB2-and RON-derived extended acceptor peptides

The single-pass membrane protein PLXNB2 (**Fig. S1b**) is a plexin family receptor for semaphorins, which mediate cell-cell interactions essential for axon guidance, cell migration, cytoskeletal dynamics and tissue organization [25, 26]. We previously identified a TMEM260-specific O-mannosylation site at T822 within the “B” β-strands of the PLXNB2 IPT1 domain [7, 13] (**Fig. 2a**). To test whether TMEM260 recognizes folded or unfolded substrates, we used a synthetic peptide encompassing the “A” and “B” β-strand segments of PLXNB2 IPT1 (residues 803-832; P^803^VITRIQPETGPLGGGIRIT^822^ILGSNLGVQ^832^) as an extended acceptor substrate model.

**Fig. 2.**
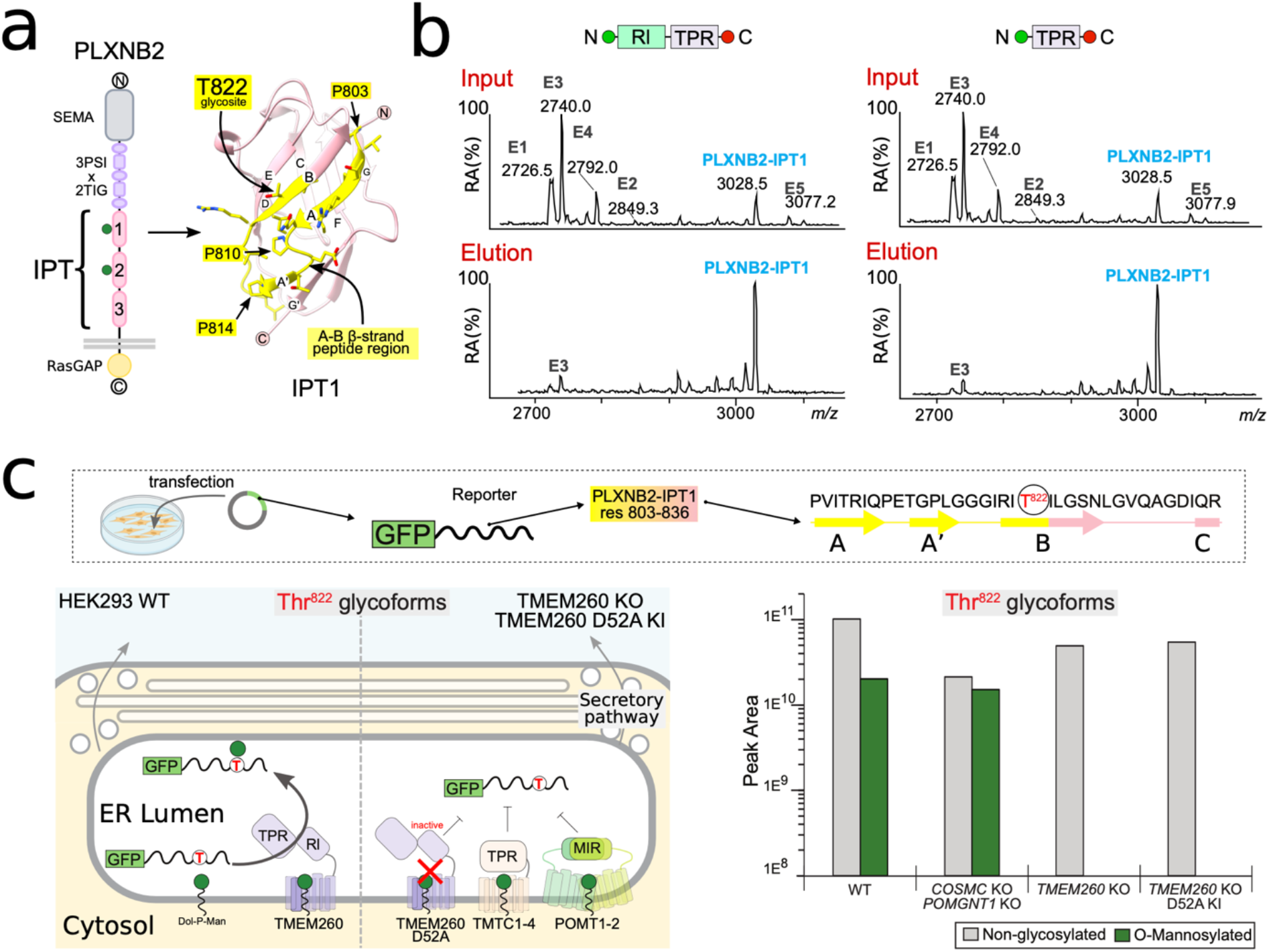
The PLXNB2-IPT1 peptide binds the luminal region of TMEM260 *in vitro* and is O-mannosylated in cells. **a**, Domain organization of the TMEM260 substrate PLXNB2, highlighting IPT domains containing reported O-mannosylation sites. The AlphaFold-predicted structure of the PLXNB2-IPT1 domain is shown in pink with annotated β-strands. The T822 glycosylation site and characteristic IPT-domain prolines within the A-B strand region are indicated. **b**, MALDI-TOF analysis of peptide pulldowns using beads decorated with TMEM260 luminal truncations (Rl-TPR or TPR-only). The TPR region is necessary and sufficient for recruitment of the PLXNB2-IPT1 peptide. **c**, Schematic (top left) of the sfGFP-PLXNB2-IPT1 reporter assay used to assess TMEM260-catalyzed glycosylation in cells. Glycoproteomic analysis of T822 glycosite shows robust O-mannosylation in WT cells and in COSMC/POMGNT1 KO “SimpleCells” used as glycoengineering control, whereas O-mannose transfer to the sfGFP-PLXNB2-IPT1 reporter is abolished in HEK293^TMEM260 KO^ and HEK293^TMEM260 KO D52A KI^ cells (right). Model (bottom left) summarizing the experimental findings. In WT cells, the reporter enters the ER lumen, where the extended C-terminal peptide is recruited by TMEM260 and the T822 is O-mannosylated. In the absence of TMEM260 activity, the ER-resident O-mannosyltransferases TMTC1–4 or POMT1/2 enzymes do not act on the sfGFP-PLXNB2-IPT1 reporter.

The ER-luminal region of TMEM260 is predicted to mediate acceptor recognition through its Rl and TPR domains [13]. Rl-TPR and TPR-only fragments (**Fig. S9a-b**) were expressed as superfolder GFP (sfGFP)-mCherry fusions [27], immobilized on agarose beads, and incubated with the PLXNB2-IPT1 peptide (**Fig. S9c; Table S3**). As controls, we used 25-mer peptides from human E-cadherin (CDH1), a substrate of the related TMTC1-4 family of GT-C O-mannosyltransferases [28]. The PLXNB2-IPT1 peptide bound efficiently to both Rl-TPR and TPR-only constructs, whereas CDH1 peptides did not (**Fig. 2b; Fig. S9d-e**). These results show that the luminal region of TMEM260, and in particular the TPR domain, is sufficient for specific acceptor recognition.

We next designed a cellular reporter assay to test whether TMEM260 catalyzes O-mannosylation of extended peptides derived from its physiological substrates PLXNB2 and RON [13]. Peptides from PLXNB2-IPT1 or RON-IPT2 were fused to sfGFP, preceded by an ER signal sequence, and transiently expressed in HEK293 wild-type (HEK293^WT^), glycoengineered TMEM260 knockout (HEK293^TMEM260 KO^) cells and HEK293^TMEM260 KO^ cells complemented with the catalytically inactive TMEM260-D52A variant (HEK293^TMEM260 KO D52A KI^) (**Fig. S10a-b**). COSMC/POMGNT1 KO “SimpleCells” served as a glycoengineering control. Glycoproteomics showed robust O-mannosylation at T822 for PLXNB2-IPT1 (**Fig. 2c**) and T703 for RON-IPT2 (**Figs. S10-S11**) in HEK293^WT^ cells. In contrast, O-mannosylation was abolished in HEK293^TMEM260 KO^ and HEK293^TMEM260 KO D52A KI^ cells, demonstrating strict dependence on catalytically active TMEM260 rather than TMTC1–4 or POMT1/2 enzymes. In TMEM260-deficient cells, T822 instead acquired sialylated Gal-GalNAc O-glycans, consistent with Golgi-mediated modification during secretion (**Figs. S10-S11**). These results show that TMEM260 specifically installs O-Man on defined sites of unfolded/extended peptide substrates in the ER lumen.

### Structure of TMEM260 bound to Dol-P-Man and PLXNB2-IPT1 peptide

To capture a TMEM260-acceptor complex, we used a fluorescently labelled truncated peptide (P^803^VITRIQPETGPLGGGIRITI^823^; dansyl-PLXNB2-IPT1) bearing one residue beyond the T822 glycosite. The cryo-EM structure of TMEM260 incubated with dansyl-PLXNB2-IPT1 was determined at 3.1 Å nominal resolution (**Fig. 3a; Table S4; Figs. S3c**,**S4c**). The map resolves the peptide with T822 positioned in the active site (**Fig. 3b, PDB 9RQM**) and reveals clear adjacent density for Dol-P-Man (**Fig. 3a-b; Fig. S12**), confirming donor carryover from cells. The structure represents the ternary complex before transfer of the sugar.

**Fig. 3.**
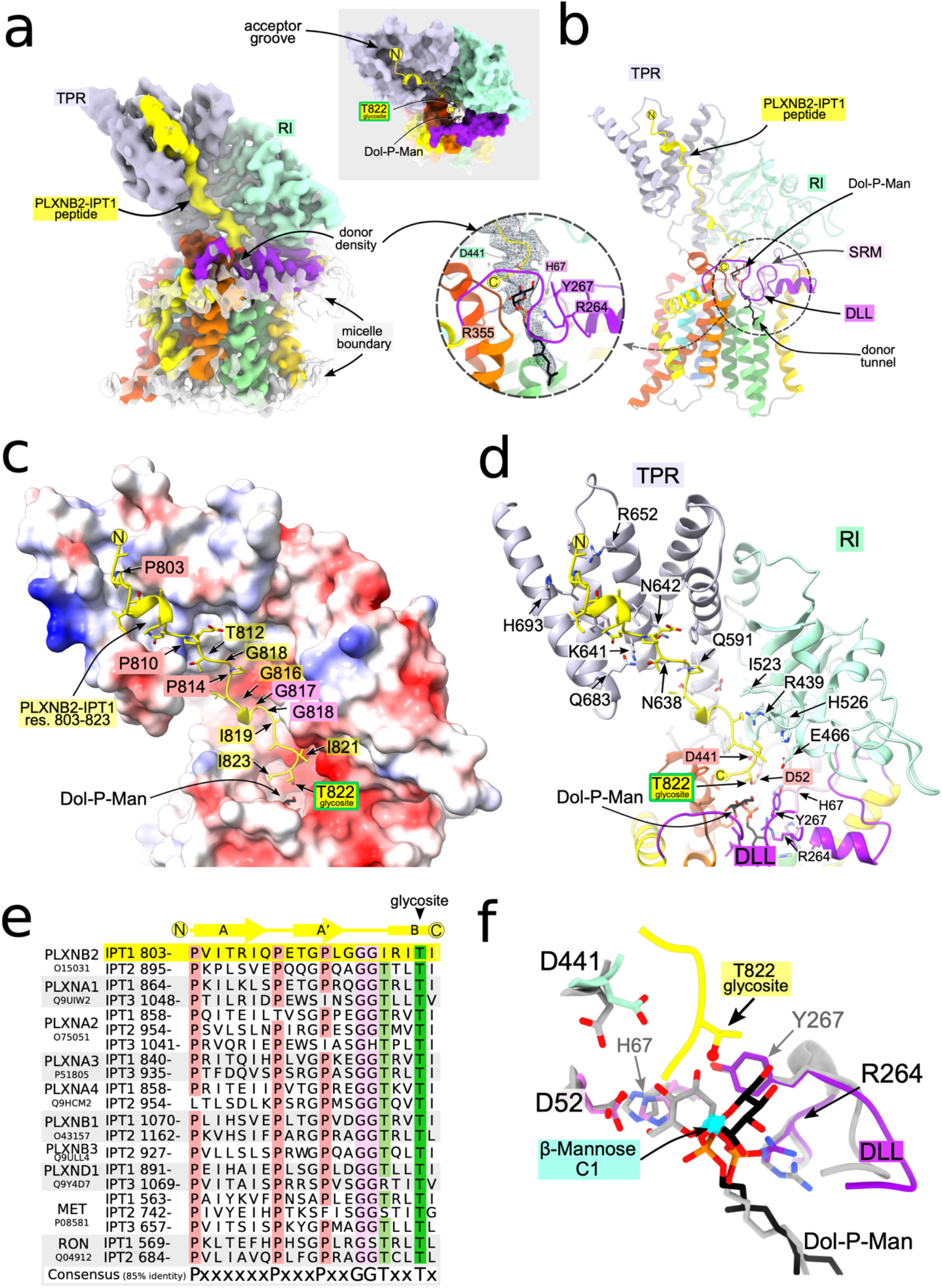
Structure of TMEM260 bond to Dol-P-Man and a PLXNB2-IPT1 peptide. **a**, Cryo-EM density map of TMEM260 in complex with the native donor Dol-P-Man and a PLXNB2-IPT1-derived acceptor peptide, determined at 3.1 Å resolution. Major structural regions are indicated and colored consistently throughout the figure. The top-right panel shows a view from the ER lumen, perpendicular to the membrane, with the acceptor peptide shown as a ribbon, the T822 glycosylation site highlighted and Dol-P-Man shown as sticks. This orientation emphasizes insertion of the N terminus into the finger region and positioning of the glycosylation site within the catalytic pocket. **b**, Front view of the corresponding atomic model in ribbon representation. Close-up (dashed circle) highlights the Dol-P-Man density and fitted model. **c**, Close-up of the ER-luminal region of TMEM260 shown in surface representation and colored by electrostatic potential, with the bound peptide as a ribbon and side chains as sticks. Dol-P-Man is shown as sticks and the T822 glycosylation site is indicated. **d**, Same view as in **c**, shown in ribbon representation to highlight specific interactions between TMEM260 residues and the acceptor peptide. The T822 glycosylation site is indicated. **e**, Sequence alignment of the PLXNB2-IPT1 peptide with reported O-mannosylated IPT domains from plexin, cMET and RON receptors, highlighting conserved residues proposed to define a TMEM260-specific acceptor sequon. **f**, Close-up of the active-site environment showing superposition of Dol-P-Man in the binary complex (all in grey including Dol-P-Man) and in the ternary complex (all in color with acceptor peptide in yellow and Dol-P-Man in black) with the PLXNB2-IPT1 peptide. In the ternary complex, the β-mannose moiety undergoes an approximately 60° rotation, indicating a peptide-induced reorientation of the donor sugar.

The ER-luminal region of TMEM260 grips the peptide (residues. 803–823) in an antiparallel orientation relative to the enzyme (**Fig. 3b**). The substrate spans a continuous groove that begins at the base of the fingers (TPR domain), runs along the palm (Rl domain), and terminates near the wrist at the transmembrane-to-luminal junction, where the catalytic pocket lies. The peptide adopts an extended conformation stabilized by regularly spaced prolines and short one-turn α-helices near its N-terminus and within the glycine triplet preceding T822. Because this peptide contains only one residue beyond T822, the structure does not capture the full C-terminal loading trajectory. Nonetheless, the extended configuration, together with our glycoproteomic data (**Fig. 2**), supports TMEM260 can modify unfolded peptide substrates, consistent with co-translational O-mannosylation.

The concave inner surface of the fingers is enriched in basic residues that engage the N-terminal half of the peptide (**Fig. 3c-d**). Residues 803–818 contact R652, H693, E683, N638, and Q591, with additional hydrogen bonds contributed by K641 and N642 to T812. Toward the palm (Rl domain), the groove becomes more neutral and culminates in a negatively charged pocket that defines the catalytic crevice. Here, the palm clasps the C-terminal segment: I819 contacts R439, G440, P443, and I523, while I821 contacts E466. T822 lies adjacent to D441 of the palm, suggesting a direct role in catalysis. The peptide terminus is anchored by I823 interactions with F349 and V352.

The binding interface defines a candidate acceptor sequon (**Fig. 3e**). By analogy, OST targets an N-x-S/T sequon in N-glycosylation [29], whereas C-mannosylation targets a W-x-x-W/C sequon [30]. On the basis of the structure and IPT domain sequence alignments, including the PLXNB2-IPT1 and RON-IPT2 substrates (**Fig. 3e; Table S1; Fig. S13**), we propose that TMEM260 recognizes a longer P-x-x-x-x-x-P-x-x-x-G-P-x-x-G/A-G-G-x-x-T/S signature that incorporates conserved residues critical for IPT-domain folding [31–33].

In the ternary complex, the Dol-P-Man sugar lies at the membrane-lumen interface beneath the acceptor groove (**Fig. 3b; Fig. S12**). The β-mannose remains in a ^4^C_1_ chair conformation, however, upon acceptor binding the sugar rotates by approximately 60°, orienting the anomeric carbon toward T822 for nucleophilic attack (**Fig. 3f**). The dolichyl chain extends into the donor tunnel below a DLL that is more disordered than in the binary complexes (**Fig. S6**).

### A catalytic mechanism coupling the transmembrane and luminal domains

TMEM260 is predicted to transfer α-mannose from Dol-P-Man with anomeric inversion (**Fig. S1a**). Inverting GTs typically catalyze a displacement reaction, in which the acceptor hydroxyl attacks the donor anomeric carbon after activation by a general base [22, 23]. The structures nominate candidate residues involved in catalysis and donor binding (**Figs. S6**,**S12**,**S14e**). We tested the effect of substituting these residues using an *in-cellulo* O-mannosylation assay [13] in HEK293^KO^ (lacking TMEM260) cells expressing a cMET ectodomain reporter (ecto-cMET; **Fig. S14a**). Reintroduction of WT TMEM260 restored >75% site occupancy across the ecto-cMET IPT1-3 domains, whereas expression of TMTC3, a related GT-C O-mannosyltransferase for cadherins [28], did not (**Fig. S14b**).

The highly conserved D441 in the Rl domain lies adjacent to the glycosite T822 and is the only acidic residue positioned to act as a general base. Consistently, the D441A substitution abolished eco-cMET glycosylation, identifying D441 as the catalytic base that activates the acceptor hydroxyl (**Fig. S14c, S15**). D52, a conserved acidic residue in the SRM of the transmembrane GT-C domain, was also essential (D52A inactive; D52E ~20% activity). Although the equivalent residue has been proposed as the catalytic base in other GT-Cs [34], in the ternary complex D52 is displaced from T822 but lies ~6 Å from the mannose anomeric carbon (**Fig. 3f**), supporting a role in productive donor binding rather than direct catalysis. Consistently, mutation of H67, another SMR residue proximal to the donor, also abolished activity. Substitutions R264A and Y267A in the DLL similarly eliminated activity, consistent with their roles in phosphate and sugar binding. Whereas, the substitution R355A in TMH 10 reduced activity to ~20%. Disruption of the asparagine ladder within the TPR domain (N602A, N638A, N642A and N680A) reduced activity to ~10%, highlighting the importance of the fingers in acceptor recruitment. Other substitutions within the ER-retention motif (R21A, R22A/S) or luminal regions (V494A, E582Q, H562A, E582A) had minimal effects, whereas C522S and C544S impaired protein stability (**Figs. S14d**,**S15**).

Together, the structures and mutagenesis support a coordinated mechanism in which the GT-C wrist binds and orients Dol-P-Man, the TPR fingers engage an extended acceptor segment, and the Rl palm positions the glycosylation site for catalysis. D441 deprotonates the acceptor hydroxyl to trigger nucleophilic attack on the anomeric carbon, while D52 stabilizes donor alignment to facilitate inversion (**Fig. S16a**). This division of labor between transmembrane donor binding and ER-luminal acceptor engagement distinguishes TMEM260 from other GT-C enzymes.

### Discussion

The importance of protein O-mannosylation for mammalian development is underscored by the spectrum of congenital muscular dystrophies, neurodevelopmental syndromes, and severe cardiac defects linked to loss-of-function mutations in the POMT1/2, TMTC1–4 and TMEM260 enzyme families [8, 15, 35, 36]. Yet the structural basis of the initial transfer of mannose to serine or threonine residues has remained unknown, limiting mechanistic interpretation of how O-Man is targeted and how these biosynthetic pathways fail in CDG.

Our cryo-EM structures define the molecular architecture of a mammalian O-mannosyltransferase. Human TMEM260 combines a transmembrane GT-C that binds Dol-P-Man with an ER-luminal substrate-recognition module that engages extended polypeptides. In this configuration, the TPR domain fingers clasp the unfolded “A” β-strand segment of the IPT domain, while the Rl domain palm positions the “B” β-strand glycosylation site within a catalytic pocket adjacent to D441. The structures further reveal donor rearrangement upon acceptor binding, aligning the anomeric center for transfer.

Akin to the C-mannosylation of W-X-X-W/C motifs in thrombospondin type domains by DPY19L1-4 [37], TMEM260 catalyzed O-Man transfer to IPT domains represents a terminal, non-elongating glycosylation event along the secretory pathway. Unlike α-dystroglycan O-Man glycans, which are elaborated into complex polysaccharide chains, O-Man on IPT domains remains unextended. Previous global O-Man glycoproteomics suggested strict specificity for IPT domains [13], leading to the hypothesis that TMEM260 acts on folded substrates. By contrast, we show that TMEM260 can selectively O-mannosylate an IPT-derived acceptor peptide containing a conserved sequon in an extended conformation, consistent with co-translational glycosylation of nascent chains before or during IPT-domain folding in the ER (**Fig. 4**). This mechanism provides a structural rationale for the receptor maturation and trafficking defects observed in TMEM260-deficient cells [13] and supports a role for TMEM260-mediated O-Man transfer in promoting IPT-domain folding and ER export of plexin, cMET and RON receptors.

**Fig. 4.**
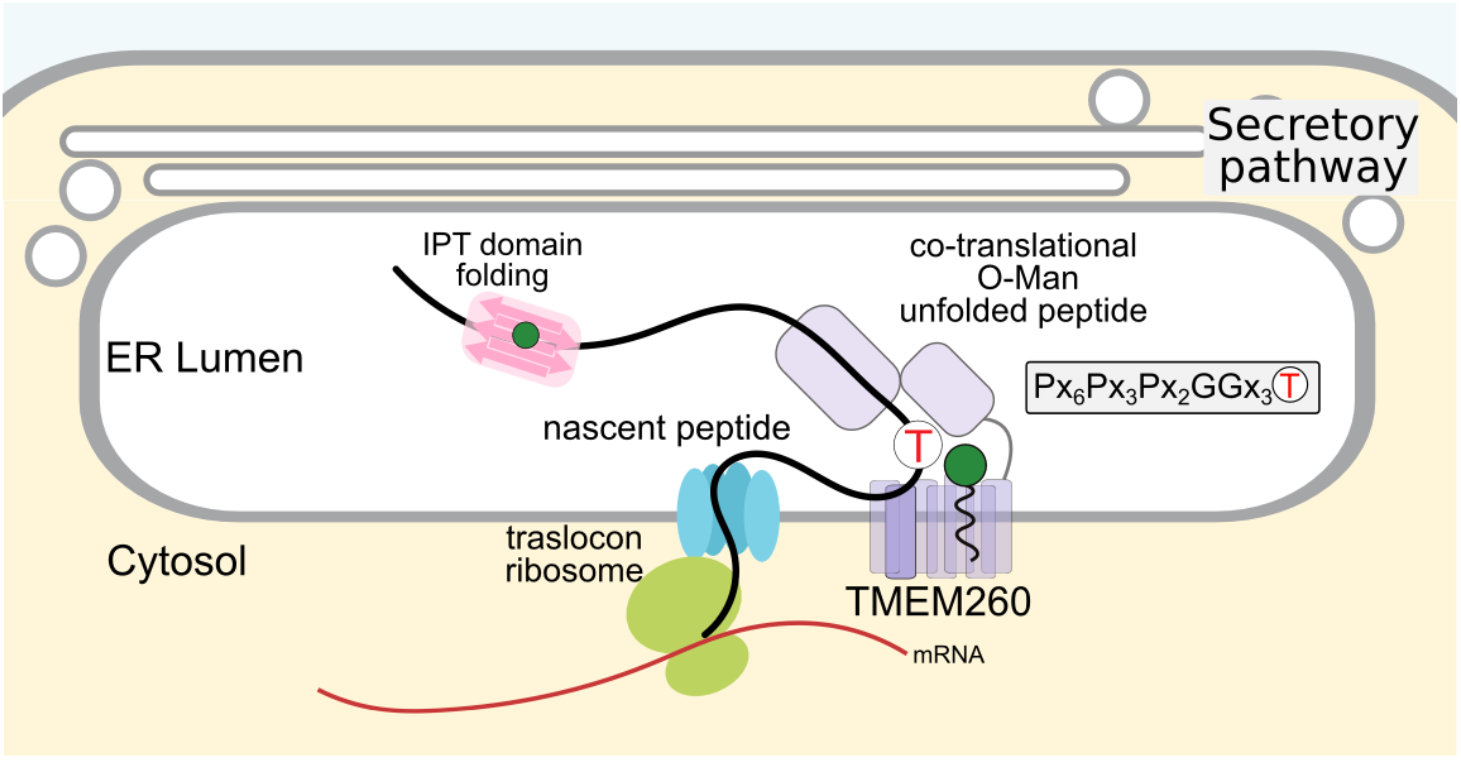
Proposed co-translational O-Mannosylation pathway mediated by TMEM260. **a**, Schematic of the proposed co-translational O-Man pathway. TMEM260 binds unfolded nascent polypeptides containing IPT domains in the ER. Exposure of a conserved sequon positions the glycosylation site within the active site for O-mannose transfer. Following release from TMEM260, the IPT domain folds and the protein proceeds through the secretory pathway to its final destination.

Finally, the structures provide a framework to interpret SHDRA-associated TMEM260 variants (**Table S2; Fig. S17**) [13–21]. Early frameshifts such as V65Afs*[33] and E55Ffs*[24], which disrupt fetal development [15], are predicted to eliminate the GT-C core and abolish O-Man transfer. Whereas truncations removing the luminal domains, Q465*[14], Y470* [15], and L447Vfs [17, 20], would disrupt acceptor binding and produce severe combined cardiac and CNS phenotypes. The recurrent W539Cfs* truncation [17], which accounts for ~26% of Japanese cases of PTA [19], disrupts the final βD–E hairpin and αT helix, likely destabilizing the luminal hand and affecting O-Man transfer. By contrast, truncations confined to distal TPR helices, such as Y567Tfs* [14], would be expected to impair recruitment with less impact on catalysis. Missense variants such as C98Y and C453R, compatible with survival into childhood, do not directly contact catalytic residues. C98Y lies beneath the active site near the E144-containing loop, whereas C453R introduces steric clashes in the palm near D52. As such these variants retain enzymatic activity [13] but likely destabilize local structure or donor positioning.

In summary, we define the architecture and mechanism of a newly identified O-Man glycosylation pathway involved in receptor folding, signalling and developmental homeostasis in mammals. TMEM260 employs a split active site in which donor binding in the transmembrane wrist is coupled to acceptor engagement and catalysis in the ER-luminal hand. This mechanism explains TMEM260’s specificity for IPT domains, defines the catalytic base responsible for inversion, and provides a structural foundation for understanding TMEM260-associated CDGs and their developmental pathology.

## Lead contact

Further information and requests for resources and reagents should be directed to and will be fulfilled by the lead contact, Iban Ubarretxena-Belandia.

## Materials availability

The glycoengineered HEK293 cells and constructs described in this study are freely available upon request from the corresponding authors. No other unique reagents were generated in this study.

## Data availability

The mass spectrometry (glyco)proteomics data have been deposited in the ProteomeXchange Consortium via the PRIDE [38] partner repository with the dataset identifier PXD072761. Single-particle cryo-EM density maps and atomic models have been deposited at the EMDB (density maps) and PDB (atomic models) and are available as follows: TMEM260 complex with Far-P-Man (C1 symmetry; EMDB 54178, PDB 9RQL), TMEM260 complex with Dol-P-Man (C1 symmetry; EMDB 54180, PDB 9RQN), and TMEM260 complex with Dol-P-Man and PLXNB2-IPT1 peptide (C1 symmetry; EMDB 54179, PDB 9RQM). The raw image datasets have been deposited at the public repository EMPIAR as follows: TMEM260:Far-P-Man (EMPIAR-13166), TMEM260:Dol-P-Man (EMPIAR-13165), and TMEM260:Dol-P-Man: PLXNB2-IPT1 (EMPIAR-13167).

## Acknowledgments

We are grateful to colleagues at the Copenhagen Center for Glycocalyx Research for fruitful discussions. I.U.-B. acknowledges funding from the Spanish Ministry of Science, Innovation and Universities (MICIU), the Spanish Research Agency (AEI) and the European Regional Development Fund (MICIU / AEI / 10.13039/501100011033 / FEDER, UE) through grant PID2022-143177NB-I00. J.P.L.-A. acknowledges financial support from a PTA contract granted by the MICIU / AEI. J.O.C. acknowledges financial support from MICIU (Recovery, Transformation and Resilience Plan) and the Basque Government “Biotechnology Complementary Plan Applied to Health” with funding from European Union NextGenerationEU (PRTR-C17.I1; PRTR-C17.I01.P01.S13). A.H. acknowledges financial support from the Carlsberg Foundation (CF21-0655), a research grant (00025438) from VILLUM FONDEN, the Novo Nordisk Foundation (NNF25OC0100678), and the Danish National Research Foundation Grants DNRF107 and DNRF196. Single-particle cryo-EM data collection was performed at the Basque Resource for Electron Microscopy located at Instituto Biofisika (UPV/EHU, CSIC), supported by the Department of Science, Universities and Innovation and the Innovation Fund of the Basque Government, by Fundación Biofísica Bizkaia and with additional support from MICIU (Recovery, Transformation and Resilience Plan) and the Basque Government “Biotechnology Complementary Plan Applied to Health” with funding from European Union NextGenerationEU (PRTR-C17.I1; PRTR-C17.I01.P01.S13; AAAA_ACG_AY_2539/22_05).

## Author contributions

Conceptualization: J.O.C., L.P., J.F.-M., A.H., I.U.-B.; Methodology: J.O.C., L.P., J.P.L.-A., I.T., J.F.-M., A.H., I.U.-B.; Investigation: J.O.C., L.P., B.O.-L., S.A., J.P.L.-A., I.T., A.H.; Resources: L.P., B.O.-L., S.Y.V., J.P.L.-A., I.T., A.H., I.U.-B.; Data curation: J.O.C., J.P.L.-A., I.T., A.H. I.U.-B.; Formal analysis: J.O.C., L.P., J.P.L.-A., I.T., H.J.J., A.H.; Visualization: J.O.C., H.J.J.; Supervision: A.H., I.U.-B.; Project administration: A.H., I.U.-B.; Funding acquisition: A.H., I.U.-B.; Writing of original draft: J.O.C., L.P., A.H. I.U.-B.; Review & editing: all authors.

## Declaration of interest

Lorenzo Povolo is currently employed by Novo Nordisk A/S; the company had no influence on the contents of this work. The other authors declare that they have no competing interests.

## Methods

### Design of the luminal domains constructs and stable cell-line generation

TMEM260 luminal domain constructs were designed as modular fusion proteins comprising, from N-to C-terminus: a CD33 (amino acids 1–16, UniProt P20138) secretion signal, a 12×His tag, eGFP, the TMEM260 Rl-TPR or TPR-only, mCherry2C (Fluorescent Protein Database; https://www.fpbase.org/), and a 3×FLAG tag. Constructs were synthesized (GenScript), subcloned into pIRES-pPuro3 (gift from Yoshiki Narimatsu, Copenhagen Center for Glycomics), and transfected into HEK293 cells using Lipofectamine 3000 (Thermo Fisher). Transfected adherent cells were maintained in DMEM supplemented with 10% fetal bovine serum (Gibco), 1% GlutaMAX (Gibco), and 1µg/mL puromycin (Gibco) until stable, puromycin-resistant, and high-expressing populations were established. eGFP fluorescence was monitored throughout selection and maintenance. Stable adherent populations were gradually adapted to suspension culture in FreeStyle™ F17 Expression Medium (Gibco) supplemented with 0.1% Poloxamer-188 (Sigma-Aldrich), 2% GlutaMAX, and 1µg/mL puromycin. Cell clumping was minimized by gentle pipetting during adaptation; debris was removed by mild centrifugation and medium replenishment. Suspension cultures were maintained at 37°C and 5% CO_2_ with shaking until viability exceeded ~90% and eGFP expression was monitored periodically. For recombinant protein production, suspension-adapted cultures were grown and conditioned medium was harvested no less than 5 days after medium replenishment to allow accumulation of secreted target proteins. Supernatants were clarified by centrifugation (3,000×g, 15min, 4°C) and filtered through 0.22-µm membranes.

### Affinity purification of luminal domains-fusion proteins

Luminal domain fusion proteins were captured from clarified supernatants using pre-cleared Ni– NTA agarose (Qiagen; 1mL slurry per 100 mL supernatant). Supernatants were adjusted to binding buffer (50mM sodium phosphate, pH 8.0, 0.5 M NaCl, 15 mM imidazole) and incubated with resin overnight at 4°C with gentle rotation. Resin was transferred to polypropylene columns (Thermo Fisher) and washed sequentially with 4 column volumes (CV) of binding buffer and 4CV of stringent wash buffer (50mM sodium phosphate, pH 8.0,1.5 M NaCl, 20 mM imidazole). Proteins were eluted in 50 mM sodium phosphate, pH 8.0, 50 mM NaCl, 450 mM imidazole and buffer-exchanged into 50 mM HEPES, pH 7.4, 50 mM NaCl using Zeba spin columns (Thermo Fisher). Final purification was performed by size-exclusion chromatography on a Superdex 75 Increase column (Cytiva) equilibrated in 50mM HEPES, pH 7.4, 50 mM NaCl. Protein purity and integrity were assessed by SDS–PAGE and by eGFP/mCherry fluorescence.

Purified luminal fusion proteins were covalently coupled to NHS-activated agarose (Pierce NHS-Activated Agarose, Thermo Fisher) according to the manufacturer’s protocol. Briefly, dry resin was swollen in amine-free coupling buffer (50mM HEPES, pH 7.4, 50 mM NaCl), mixed with protein in the same buffer, and incubated for 1–2 h at room temperature with end-over-end rotation. Unreacted NHS groups were quenched with 1M Tris for 15-20 min, and conjugated resin was washed extensively with coupling buffer. The Rl-TPR and TPR-only conjugated resins were used for peptide enrichment as described below.

### Peptide design and enrichment

A PLXNB2-derived peptide was designed from sequence alignments of human IPT domains using the UniProt alignment tools; conserved regions were identified and a representative sequence selected for synthesis (Schafer-N, Copenhagen). Control peptides were designed from human CADH1 sequence and acquired similarly. For enrichment assays, peptides were incubated with Rl-TPR and TPR-only conjugated resins for 2h at room temperature in 50mM HEPES, pH 7.4, 50mM NaCl. Beads were washed three times with the same buffer and bound peptides were eluted with 2M NaCl. Eluates were desalted using C18 Sep-Pak cartridges (Waters) following the manufacturer’s instructions and as described [7] and peptides were eluted in 50% acetonitrile (ACN) with 0.1% formic acid (FA), and prepared for MALDI-TOF analysis.

### MALDI-TOF sample preparation and acquisition

For MALDI-TOF, 1µL of eluted peptide solution was spotted onto an MTP AnchorChip™ 384 BC target plate (Bruker Daltonik GmbH, Bremen, Germany) and mixed on-plate with 1µL super-DHB (sDHB; 5mg/mL in 50% ACN, 0.1% FA) matrix. Samples were air-dried at room temperature. Spectra were acquired on a Bruker Autoflex MALDI-TOF/TOF mass spectrometer using FlexControl 3.4 in positive-ion reflector mode (m/z 900-4500), accumulating up to 10000 laser shots per spectrum at 100 Hz. Raw spectra were processed in FlexAnalysis 5.1 using default baseline subtraction, smoothing and peak-picking parameters.

### Plasmid construction and TMEM260 variant generation

For variant cloning, the pIRES-pPuro vector was modified by PCR to introduce BamHI and NotI restriction sites using oligonucleotides listed in the Reagents section (Oligos 1 and 2). PCR was performed with KOD Hot Start DNA polymerase (Sigma-Aldrich), followed by DpnI (NEB) digestion to remove template plasmid. Products were gel-purified, assembled using In-Fusion (TaKaRa) and transformed into Stellar competent cells (TaKaRa). The modified vector was digested with BamHI and NotI (NEB, high-fidelity enzymes), and TMEM260 WT inserts (TMEM260-3×FLAG on EPB71 [28]) were cloned by In-Fusion-assisted ligation. Site-directed mutagenesis was performed using variant-specific primers with the same KOD/DpnI workflow. For the asparagine-ladder mutant, the template was linearized by PCR and assembled by In-Fusion with a synthetic DNA cassette (IDT) encoding the desired mutations. All constructs were validated by Sanger sequencing (Eurofins Genomics).

### Plasmid construction for sfGFP-PLXNB2-IPT1 and sfGFP-RON-IPT2 peptide reporters

Reporter sfGFP-PLXNB2-IPT1 and sfGFP-RON-IPT2 constructs comprised sfGFP fused at its with C-terminus with the target region of PLXNB2 (803-837; PVITRIQPETGPLGGGIRIT^822^ILGSNLGVQAGDIQR) or RON (684-715; PVLIAVQPLFGPRAGGTCLT^703^LEGQSLSVGTSR). The coding region for PLXNB2 (803-837) was amplified from pENTR223-PLXNB2 (Addgene plasmid #86236; gift from Roland Friedel; RRID:Addgene_86236) and RON (684-715) from pDONR223-MST1R (Addgene plasmid #23942; gift from William Hahn & David Root; RRID:Addgene_23942). Inserts were cloned by In-Fusion (TaKaRa) into a backbone generated by inverse PCR from the TMEM260 luminal-domain expression plasmid series (maintaining pIRES-Puro3-sfGFP-polyHis-TEV) using 15-bp homology extensions. Primer sequences used for amplification were: for inverse PCR of template; forward primer 5’-TAAGCGGCCGCATAGATAACTGA-3’, reverse primer 5’-TGATTGGAAGTAAAGATTCTCCCCA-3’. Amplification of PLXNB2 (803–837); forward primer 5’-CTTTACTTCCAATCACCCGTCATCACCAGGATCC-3’, reverse primer 5’-CTATGCGGCCGCTTACCTCTGGATGTCCCCTGC-3’. Amplification of RON (684-715); forward primer 5’-CTTTACTTCCAATCACCAGTGCTGATAGCAGTGCAAC-3’, reverse primer 5’-CTATGCGGCCGCTTACCGGCTGGTGCCTACAGAC-3’. Reporters were assembled as ER-signal-(7×G)-(12×H)-(6×G)-sfGFP-(7×G)-(TEV site: ENLYFQ↓S)-peptide. Constructs were transformed into competent *E. coli*, purified (QIAGEN) and validated by Sanger sequencing (Eurofins Genomics).

### ecto-cMET reporter assay

A donor cell line carrying TMEM260 knockout and an ecto-cMET reporter (ectodomain of cMET) knock-in at the AAVS1 locus was used for variant expression [13, 28]. Cells were transfected with TMEM260 variant plasmids using Lipofectamine LTX with Plus reagent (Invitrogen) and maintained in DMEM with 10% FBS, 2mM GlutaMAX and puromycin at the minimal concentration required for complete lethality (determined previously by titration at 0.75ug/mL). Cells were plated on poly-L-lysine-coated 10-cm dishes (0.1mg/mL; Sigma) and grown to >80% confluence. Medium was replaced with serum-free OptiMEM (Thermo Fisher), and cells were cultured for 5 days without replenishment to accumulate secreted ecto-cMET reporter in the medium. Conditioned media were clarified (3,000×g, 15min, 4°C), filtered (0.22 µm) and immunopurified with anti-FLAG M2 magnetic beads (Sigma-Aldrich). Binding was performed overnight at 4°C with gentle rotation. Beads were washed three times with 50 mM HEPES, pH7.4, 150mM NaCl, and bound proteins were eluted in 10% SDS. Eluates were processed on S-Trap Micro columns (ProtiFi) following the Micro High Recovery protocol with ammonium bicarbonate in place of TEAB. Proteins were reduced (10mM DTT, 30min, 56°C), alkylated (20mM iodoacetamide; 30min, room temperature, in darkness), quenched with 10mM DTT, supplemented to 5% SDS, and digested overnight at 37°C with sequencing-grade modified trypsin (Promega) in a humidified chamber. Peptides were desalted on in-house packed StageTips (Empore C18,3M) prior to LC-MS/MS.

### Affinity purification of sfGFP reporters, western blotting and sample preparation for bottom-up glycoproteomics

sfGFP-PLXNB2-IPT1 (803-837) and sfGFP-RON-IPT2 (684-715) reporters were transiently expressed in WT and glycoengineered HEK293 cells [13] using Lipofectamine LTX (Thermo Fisher) according to the manufacturer’s instructions. Transfected cells were cultured under conditions described above, and 20 mL of serum-free medium was harvested 72 h post-transfection and clarified (1000 × *g*, 10 min, 4°C). Reporters were purified by Ni-NTA chromatography, as described above for luminal domain. Eluates were separated by SDS-PAGE (4-12% Bis-Tris gels) in MES buffer before transfer (100V, 60 min) to PVDF membranes and western blot analyses using HRP-conjugated GFP monoclonal antibody (GF28R; Thermo Scientific) at 1:4000 dilution in 5% skim milk, TBST buffer. For glycoproteomics, approximately 20 µg of purified reporter from each condition was digested on S-Trap micro spin columns (ProtiFi) according to the manufacturer’s protocol with 1 µg trypsin (Roche). Peptides were desalted on in-house packed StageTips (Empore C18, 3M) prior to LC-MS/MS.

### LC-MS/MS acquisition and data analysis for ecto-cMET reporter

LC-MS/MS was performed essentially as previously described [13]. Peptides were analyzed on an EASY-nLC 1000 coupled to an Orbitrap Fusion Tribrid or Fusion Lumos mass spectrometer via a nanoSpray Flex source (Thermo Fisher Scientific). Samples were separated on in-house packed analytical column (PicoFrit emitter, 75µm i.d.; New Objectives) containing Reprosil-Pure-AQ C18 (1.9µm; Dr. Maisch) and operated at 200nL/min with a 90-min gradient using solvent A (0.1% FA) and solvent B (ACN, 0.1% FA). Precursor MS1 scans (m/z 355-1700) were acquired at 120,000 resolution followed by HCD-MS/MS and ETciD-MS/MS of multiply charged precursors (z=2-6). A minimum MS1 signal threshold of 10,000-50,000 ions was used to trigger data-dependent fragmentation. MS2 spectra were acquired at 60,000 resolution. Raw files (.raw) were processed in Proteome Discoverer 1.4 (Thermo Fisher Scientific) using the Sequest HT node and searched against a FASTA containing full-length cMET (P08581, MET_HUMAN; UniProt download 2024-07-01) with parameters: precursor tolerance 10ppm, fragment tolerance 0.02 Da, up to two missed tryptic cleavages (full- and semi-specific), carbamidomethylation (C; 57.02146 Da) set as fixed, oxidation (M; 15.9949 Da) and Hex (162.0528 Da) on Ser/Thr as variable modifications. Peptide confidence was assessed with the Target-Decoy PSM Validator and filtered at p<0.01 for high-confidence identifications. All spectral matches for glycopeptides were inspected manually. Identified peptides and glycopeptides were further processed in Skyline [39] (v25.1.0.142; MacCoss Lab Software) for estimation of O-Man site occupancy. Mass spectrometry data is based on single-shot analyses of individual ecto-cMET samples which precludes statistical analyses.

### LC–MS/MS acquisition and data analysis for sfGFP reporter proteins

sfGFP reporters were analyzed as above with minor modifications. Peptides were separated using a 60-min gradient on an EASY-nLC 1000 coupled to a Fusion Lumos Tribrid mass spectrometer (Thermo Fisher Scientific). Multiply charged precursors were fragmented by Collision-Induced Dissociation (CID), Higher-Energy Collision Dissociation (HCD), and hybrid Electron-Transfer Collision-Induced Dissociation (ETciD). Raw data were processed using FragPipe (v23.0) and MSFragger (v4.3) with the “glyco-O-hybrid” workflow [40] against a FASTA containing the sfGFP reporter sequences supplemented with decoys and common contaminants. Glycopeptide spectra were manually inspected. Label-free quantification and AUC calculations were performed in Skyline [39] (v25.1.0.142; MacCoss Lab Software). Mass spectrometry data is based on single-shot analyses of individual sfGFP reporter samples which precludes statistical analyses.

### Cell culture, transient expression and protein purification for cryo-EM

Expi293F cells (Thermo Fisher, A14527) were maintained and transfected in suspension in Expi293™ Expression Medium (Thermo Fisher, A1435102) in an orbital incubator at 37°C, 8% CO_2_ and 135rpm. For large-scale TMEM260 expression, cultures were adjusted to 2×10^6 cells/mL the day before transfection and transfected at ~3×10^6 cells/mL with TMEM260-3×FLAG on EPB71 plasmid [28] using PEIPro (Polyplus) at a 1:1 (w/w) DNA:PEI ratio (1.5µg DNA/mL culture). After 48h, cells were harvested (3,000 rpm, 5min) and pellets were flash-frozen. Frozen pellets were resuspended to 200 mg cells/mL in resuspension buffer (TBS with 150 mM NaCl; 50 mM Tris-HCl, pH 7.4, 5% glycerol, 2 mM MgCl_2_, 0.5 mM PMSF, 1mM benzamidine and 25 µg/mL DNase I) and homogenized with a Dounce homogenizer. DDM was added to 1.5% (w/v) and lysates were solubilized for 1h at 4°C with rotation at 15rpm. Insoluble material was removed by ultracentrifugation (58,000×g, 40min, Beckman Optima LX-100, 70Ti rotor). Clarified lysate was incubated with anti-FLAG G1 (1mL, pre-equilibrated in TBS, 2mM MgCl_2_, 0.025% DDM) for 1h at 4°C with gentle agitation. The mixture was applied to a gravity flow column, washed (15CV) and elute (5CV) with wash buffer containing 200 µg/mL FLAG peptide. Eluted fractions were concentrated to 550µL (Amicon 100-kDa cutoff) and further purified by size-exclusion chromatography on a Superdex 200 10/300 column equilibrated in TBS with 2mM MgCl_2_ and 0.025% DDM. Monodisperse peak fractions were pooled for cryo-EM.

### Cryo-EM grid preparation and data collection

Freshly purified TMEM260 was used for vitrification. For ternary complex formation, TMEM260 at 2 mg/mL was mixed with dansylated peptide (3 mM in DMSO) to 0.3 mM final concentration and incubated on ice for 15 min. For the donor-analog complex, TMEM260 at 1mg/mL was incubated with Far-P-Man stock (50 mM in DMSO) at 0.25 mM final concentration for 1h at 4°, and then concentrated to 3.8 mg/mL. For the native Dol-P-Man complex we used TMEM260 at 2 mg/mL without any exogenous substrates. UltrAuFoil R1.2/1.3 300-mesh grids were glow-discharged (3 min, 10 mA, ELMO Glow Discharge System), loaded with 3 mL of sample, blotted for 4s and plunge-frozen in liquid ethane using a Vitrobot Mark IV (Thermo Fisher Scientific) at 4°C and 100% humidity. Specimens were imaged on a Krios G4 (Thermo Fisher Scientific) at 300kV equipped with a Gatan K3 and BioQuantum energy filter (20eV slit width) using EPU3 (Thermo Fisher Scientific) at 105,000× nominal magnification (0.8238Å/pixel, counting mode). Defocus ranged from −0.5 to −2.4 µm. Movies were recorded as 60-frame stacks with total doses of 50 e^−^/Å^2^ (ternary complex) and 60.1 e^−^/Å^2^ for Far-P-Man and Dol-P-Man binary complexes (**Table S4**).

### Cryo-EM image processing

TMEM260:Dol-P-Man, TMEM260:Far-P-Man and TMEM260:Dol-P-Man:PLXNB2-IPT1 movies (0.8238Å/pixel) were motion-corrected using patch motion correction (global and local), and CTF parameters were estimated using patch CTF estimation in CryoSPARC Live (v4.7) [41]. Particles were initially picked with a blob picker (particle diameter 70-200Å), extracted to a 320-pixel box and binned to 160 pixels for 2D classification. Representative 2D classes (low-pass filtered to 20Å) were used for template picking during data collection. Binned particles and aligned movies were exported for subsequent processing. Particles were subjected to iterative 2D classification and *ab initio* reconstruction in C1. Heterogeneous refinement was used to remove junk particles, followed by homogeneous refinement (HR) and non-uniform refinement (NUR) using the HR mask for an initial binned reconstruction that contained full luminal and transmembrane density. Particles were re-extracted unbinned, subjected to HR and NUR, and refined per-particle and global CTF refinement. TOPAZ [42] was trained on selected particles to recover missed particles for additional NUR. Reference-based motion correction was applied, followed by final HR, NUR, and local refinement using a soft mask around TMEM260 (**Table S4**).

### Model building and refinement

An AlphaFold model of TMEM260 (UniProt Q9NX78) was rigid-body fitted into cryo-EM maps in Chimera. Iterative model building and refinement were performed using PHENIX [43] real_space_refine and Coot [44] including rigid-body refinement of luminal and transmembrane regions followed by local grid searches and ADP refinement. Ligands were modeled using CIF files from the Grade Web Server. Final geometry improvements were performed in ISOLDE [45]. Model quality was assessed with MolProbity and the validate3D-PDB server (**Table S4**).

### Surface analysis and tunnel calculations

Cavity and tunnel analyses were performed in CAVER Analyst 2.0 [46] using probe radii ranging from 1.4-10 Å. The largest cavity had a volume of 14,144 Å^3^ (40,100 sampled points) and accommodated a probe radius up to 6.06 Å. For tunnel detection, a probe radius of 1.8 Å was used to exclude narrow channels. A clustering threshold of 7 Å, shell depth of 4 Å and shell radius of 3 Å were applied. Starting points were defined by atoms lining the largest cavity. Optimization used a maximum distance of 5 Å, desired radius of 10 Å, and default approximation.

### Analysis of sequon candidates

PROSITE-format patterns were derived from IPT domain alignments of reported targets. Patterns were scanned against the UniProt human reviewed proteome (query: organism_id:9606 AND reviewed:true AND proteome:UP000005640 AND is_isoform:false) using the ps_scan script from ExPASy SIB (ftp://ftp.expasy.org/databases/prosite/ps_scan/ps_scan_linux_x86_elf.tar.gz). For each PROSITE pattern, Shannon information content was calculated as previously described [47]. For a pattern of length “l”, total information “Rt” was computed as Rt=∑j=Rj, where Rj is the amount of information content for each position j calculated by the formula Rj=log2(N)−log2 (kj), where N=20 is the total number of standard amino acids, and “kj” is the count of allowed amino acids at position “j”. Specifically, “kj” values were assigned for the pattern as follows: value 1 for fixed amino acids (i.e. - P-defined residue maximum information in a position ≈4.32 bits), 20 for the wildcard “x” (any residue at position, 0 bits), and the number of amino acids in square bracket lists (e.g. 3 for [PFI], ≈2.74 bits). The total Shannon information content (Rt) for a pattern was the sum of Rj across positions.

