## Supplementary information for "Structure and mechanism of the human TMEM260 O-mannosyltransferase"

### Table of Contents

#### 1. Supplementary Tables

Table S1: Experimentally reported TMEM260 acceptors, IPT/TIG domain O-Man sites and predicted matches to candidate sequons.
Table S2. Literature-reported cohort of patients with TMEM260 mutations and congenital heart defects (CHDs).
Table S3. Reagents, oligonucleotides and peptides.
Table S4. Cryo-EM data collection, refinement and validation statistics.

#### 2. Supplementary Figures

Fig. S1. O-mannosyltransferase reaction and TMEM260 substrate specificity.
Fig. S2. TMEM260-3×FLAG construct, purification and donor structures.
Fig. S3. Cryo-EM data processing.
Fig. S4. Local map resolution.

Fig. S5. The transmembrane region of the TMEM260:Far-P-Man binary complex.
Fig. S6. Active site architecture in TMEM260 binary complexes.
Fig. S7. Structural comparisons of TMEM260 with representative glycosyltransferases.
Fig. S8. Conservation analysis of TMEM260 across species.
Fig. S9. PLXNB2-IPT1 pull-down by the ER-luminal domains of TMEM260.
Fig. S10. TMEM260-specific O-mannosylation of Plexin-B2 and RON receptor peptides fused to GFP-reporters.
Fig. S11. Annotated MS/MS spectra of tryptic glycopeptides from sfGFP reporters expressed in glycoengineered HEK293 cell lines.
Fig. S12. Active site architecture in the TMEM260 ternary complex.
Fig. S13. Analysis of candidate TMEM260 sequons.
Fig. S14. O-mannosylation activity of TMEM260 variants.
Fig. S15. O-mannosylation activity of TMEM260 variants in HEK293 cells.
Fig. S16. Proposed catalytic mechanism of TMEM260 compared with representative GT-C enzymes.
Fig. S17. Putative structural effects of patient-associated TMEM260 variants linked to congenital heart disease.

**3. Supplementary References**

### 1. Supplementary Tables

**Table S1. Experimentally reported TMEM260 acceptors, IPT/TIG domain O-Man sites and predicted matches to candidate sequons.** Immunoglobulin-like fold, Plexins, Transcription factors (IPTs) are also known as Transcription factor ImmunoGlobin (TIG) domains. The domain notation IPT/TIG in UniProt corresponds to InterPro entry IPR002909. "No IPT" or "No TIG" indicates that the corresponding IPT (SMART: SM00429) or TIG (Pfam: PF01833) annotation is not detected. † Denotes experimentally reported O-Man sites [1, 2]. Values (1)-(3) indicate matches to the candidate PROSITE patterns listed below; "+" indicates a match detected by pattern scanning but not experimentally reported. "x" indicates absence of Thr at the glycosylation-site position. "†matches/total†" indicates the number of experimentally reported sites recovered by the pattern relative to the total reported sites. "Total extra matches" indicates pattern matches not experimentally reported. "Matches on targets (‡)" reports the number of acceptor proteins containing at least one match.

<table>
<tr><td colspan="2">Acceptors<br>Targets</td><td>Other domain<br>at N-term</td><td>IPT/TIG<br>1</td><td>IPT/TIG<br>2</td><td>IPT/TIG<br>3</td><td>IPT/TIG<br>4</td></tr>
<tr><td colspan="2">PLXNA1</td><td>†</td><td>† (1)</td><td>x</td><td>† (2)</td><td>x</td></tr>
<tr><td colspan="2">PLXNA2</td><td></td><td>† (3)</td><td>† (1)</td><td>†(2)</td><td>x</td></tr>
<tr><td colspan="2">PLXNA3</td><td></td><td>† (1)</td><td>† (1)</td><td>(3+)</td><td>x "!"</td></tr>
<tr><td colspan="2">PLXNA4</td><td></td><td>† (1)</td><td>† (3)</td><td>x</td><td>x</td></tr>
<tr><td colspan="2">PLXNB1</td><td></td><td>† (1)</td><td>† (1)</td><td>x</td><td></td></tr>
<tr><td colspan="2">PLXNB2</td><td>†</td><td>† (1)</td><td>† (1)</td><td>x No TIG</td><td></td></tr>
<tr><td colspan="2">PLXNB3</td><td></td><td>(1+)</td><td>† (1)</td><td>x</td><td>x No IPT</td></tr>
<tr><td colspan="2">PLXND1</td><td></td><td>† (1)</td><td>(1+)</td><td>† (1)</td><td></td></tr>
<tr><td colspan="2">c-MET</td><td></td><td>† (1)</td><td>† (1)</td><td>† (2)</td><td></td></tr>
<tr><td colspan="2">RON</td><td></td><td>† (3)</td><td>† (1)</td><td></td><td></td></tr>
<tr><td colspan="2">TOTAL 10</td><td></td><td colspan="4">21†</td></tr>
<tr><td colspan="7">Sequon O-Man candidate PROSITE pattern</td></tr>
<tr><td></td><td colspan="3">PROSITE pattern</td><td>†matches<br>/total†</td><td>total extra<br>matches</td><td>matches on<br>targets (‡)</td></tr>
<tr><td>1</td><td colspan="3">P-x(6)-P-x(3)-P-x(2)-G-G-x(3)-T-[ILVG]</td><td>15/21</td><td>2</td><td>17/10</td></tr>
<tr><td>2</td><td colspan="3">P-x(6)-P-x(2)-[GAS]-[PFI]-x(2)-G-[GH]-x(2)-[ILV]-T-[ILVG]</td><td>18/21</td><td>2</td><td>20/10</td></tr>
<tr><td>3</td><td colspan="3">[PL]-x(6)-[PT]-x(2)-[GAS]-[PFI]-x(2)-G-[GHS]-x(2)-[ILV]-T-[ILVG]</td><td>21/21</td><td>3</td><td>24/10</td></tr>
</table>

**Table S2. Literature-reported cohort of patients with TMEM260 mutations and congenital heart defects (CHDs).** TOP: Terminus of Pregnancy. PTA: Persistent Truncus Arteriosus. VSD: Ventricular Septal Defect. ASD: Atrial Septal Defect. TVA: Tricuspid Valve Atresia. KD. Kidney Disease. DORV: Double-Outlet Right Ventricle. TOF: Tetralogy of Fallot. IAA: Interrupted Aortic Arch. PAPVR: Partial Anomalous Pulmonary Venous Return. F: family number at report. (s.r) surgery repaired. (‡) Family history includes previous reports of dead siblings, but they were not genotyped. (□) The first pregnancy terminated at 28 w. (!) Previously annotated as variant pArg115Lys.

| Cases Family | Zygosity | cDNA variants | Protein variants | Diagnostic | Age at report | ClinVar Variation ID | Reported in |
|---|---|---|---|---|---|---|---|
| 1♂1♀ F1 siblings | Het | c.344G>A (p.Arg115Lys) **(!)** c.160+2159_344+1212del | Val65Alafs*32 **(!)** Glu55Phefs*20 | TA type I VSD; PFO[1] | 1♂ TOP 24w 1♀ TOP 21w | 1319994 1319995 | [3] |
| 1♂ (‡1) | Hom | c.637-319_858-92delinsA | Glu213Phefs*92 | TA type II; VSD; IAA; PAPVR[2] | Died 53 days | | [4] |
| 1♀ (‡1) (□) | Hom | c.1336_1339 | L447Vfs*9 | TA type I VSD; PFO | 10m.o. | | [5] |
| 1♂1♀ siblings | Het | c.1393C＞T 7066-bp deletion exons 6-7 | Gln465* | ♂PTA ♀DORV | N.A | 426075 | [6] |
| 2♂1♀ F1 siblings | Hom | c.1393C>T | Gln465* | SHDRA | 2y.o. Died 2m.o. Died 6w.o. | 426075 | [7] |
| 1♂ F2 | Hom | c.1410C>A | Tyr470* | TA type I VSD; CKD | Died 5y.o. | 1992345 | [3] |
| 2♂ F3 brothers | Het | c.1393C>T c.1644delT | Pro549Leufs*46 Gln465* | TA type I (s.r) KD* | *Died 3m.o. 5y.o. | 1264353 426075 | [3] |
| 1♀ | Het | 1617del c.332dup | Trp539Cysfs*9 Thr112Hisfs*36 | | 16y.o. | | [8] |
| 1♂1♀ (‡) siblings | Hom | c.1617del | Trp539Cysfs*9 | PTA | ♀ 4y.o. ♂ 1y.o. | | [9] |
| 1♀ F.2 | Hom | c.1617del | Trp539Cysfs*9 | PTA | ♀6y.o. | | [9] |
| 1♀ F3 | Hom | c.1617del | Trp539Cysfs*9 | PTA | ♀22y.o. | | [9] |

| | | | | | | | |
|---|---|---|---|---|---|---|---|
| 1 ♂ F4 (‡) | Hom | c.1617del | Trp539Cysfs*9 | PTA | ♀2d.o. | | [9] |
| 1 ♂ F5 siblings | Hom | c.1617del | Trp539Cysfs*9 | PTA | ♀7y.o. | | [9] |
| 1♀ (‡) | Hom | c.1617del | Trp539Cysfs*9 | PTA | Died 3m.o. | | [8] |
| 1♀ | Hom | c.1617del | Trp539Cysfs*9 | PTA | 11y.o. | | [8] |
| 1 ♂ | Hom | c.1617del | Trp539Cysfs*9 | PTA | 29y.o. | | [8] |
| 1 N/A | Hom | c.1688del | Thr563Lysfs*32 | SHDRA | N/A | 974872 | [10] |
| 1♀ F2 (‡) | Hom | c.1698_1701 del | Tyr567Thrfs*27 | SHDRA | Died 1y.o. | 426076 | [7] |
| 1 ♂ F4 | Hom | c.1698_1701 del | Tyr567Thrfs*27 | TA type I (s.r) VSD; ASD; TOF;KD | 3y.o. | 426076 | [3] |
| 1 N/A | Hom | c.1698_1701 del | Tyr567Thrfs*27 | SHDRA | N/A | 426076 | [10] |
| 1 ♂ | Het | c.1858C > T c.1617delG | Gln620* Trp539Cysfs*9 | PHACE S-like | 10y.o. | | [11] |
| 1 ♂ | Het | c1960C>T c.1617del | Gln654* Trp539Cysfs*9 | | 6y.o. | | [8] |
| 2♀ (sisters) | Het | c.1744G>C c.193-2A>G | Glu582Gln | SHDRA | TOP 22w. Died 4m.o. | 1202597 1202596 | [3] |
| 1 | Het | c.344dup c.90_104dup | Leu116Alafs*32 Ala35_Val36insAla ValPheAlaAla | ASD; DA; renal dysplasia | | 3382584 3382585 | [12] |
| 1 ♂ | Hom | c.293 G>A | Cys98Tyr | SHDRA | 5y.o. | | [1] |
| 1 ♂ (‡) | Hom | | Cys453Arg | SHDRA | 8y.o. | | [1] |

**Table S3. Reagents, oligonucleotides and peptides.**

| Item | Sequence/Details | Vendor/Catalog/Notes |
|---|---|---|
| n-Dodecyl β-D-maltoside (DDM) | N/A | GLYCON Biochemicals GmbH |
| β-D-mannosyl farnesyl phosphate ammonium salt (donor analog) | N/A | Avanti Polar Lipids, LLC |
| Synthetic dansylated peptide | Dansyl-PVITRIQPETGPLGGGIRITI | GenScript USA Inc. (stock: 3 mM in DMSO) |
| PLXNB2-derived peptide | PVITRIQPETGPLGGGIRITILGSNLGVQ A | Schafer-N, Copenhagen |
| CDH1 peptide 1 | VSSNGNAVEDPMEILITVTDQNDNKPE F | Schafer-N, Copenhagen |
| CDH1 peptide 2 | KVTEPLDRERIATYTLFSHAVSSNG | Schafer-N, Copenhagen |
| CDH1 peptide 3 | QGADTPPVGVFIIERETGWLKVTEP | Schafer-N, Copenhagen |
| CDH1 peptide 4 | LVQIKSNKDKEGKVFYSITGQGADT | Schafer-N, Copenhagen |
| CDH1 peptide 5 | DWVIPPISCPENEKGPFPKNLVQIK | Schafer-N, Copenhagen |
| Oligo1_Vector_Fwd | 5′–TAAGaattcGCTAGCGCGGCCGCATAG ATAACTGATCC–3′ | TAGCopenhagen, used for BamHI/NotI introduction on pIRES-Puro |
| Oligo2_Vector_Rev | 5′–TAGCgaattCTTAAGGATCCGAGCTCG GTACCAAGC–3′ | TAGCopenhagen, used for BamHI/NotI introduction on pIRES-Puro |
| Oligo3_His67Ala_Fwd | 5′–CGAACTGGGAGTCGCTgcCCCCCCCG GCTACC–3′ | TAGCopenhagen, used for generation of His67Ala variant |
| Oligo4_His67Ala_Rev | 5′–GTAGCCGGGGGGGcAGCGACTCCC AGTTCG–3′ | TAGCopenhagen, used for generation of His67Ala variant |
| Oligo5_Arg264Ala_Fwd | 5′–ggtttccTCACCCACTTTTTAgccGAGGAG TACGGC–3′ | TAGCopenhagen, used for generation of Arg264Ala variant |

| | | |
|---|---|---|
| Oligo6_Arg264Ala_Rev | 5′–gaaaatgtGCCGTACTCCTCggcTAAAAAGTGGGtgag–3′ | TAGCopenhagen, used for generation of Arg264Ala variant |
| Oligo7_Tyr267Ala_Fwd | 5′–ACTTTTTAAGGGAGGAGgcCGGCACATTTTCTTTAGC–3′ | TAGCopenhagen, used for generation of Tyr267Ala variant |
| Oligo8_Tyr267Ala_Rev | 5′–CTAAAGAAAATGTGCCGgcCTCCTCCCTTAAAAAGTGG–3′ | TAGCopenhagen, used for generation of Tyr267Ala variant |
| Oligo9_Arg355Ala_Fwd | 5′–CATGGGAGTGGTGGAAgccTTCTGGATGCAAAGC–3′ | TAGCopenhagen, used for generation of Arg355Ala variant |
| Oligo10_Arg355Ala_Rev | 5′–GCTTTGCATCCAGAAggcTTCCACCACTCCCATG–3′ | TAGCopenhagen, used for generation of Arg355Ala variant |
| Oligo11_Asp441Ala_Fwd | 5′–CCATCATTTTACTGAGAGGCgccCTGCCCGGAAATTCTTTAAGG–3′ | TAGCopenhagen, used for generation of Asp441Ala variant |
| Oligo12_Asp441Ala_Rev | 5′–CCTTAAAGAATTTCCGGGCAGggcGCCTCTCAGTAAAATGATGG–3′ | TAGCopenhagen, used for generation of Asp441Ala variant |
| Oligo13_Glu466Ala_Fwd | 5′–CTTTAGTGGACCAAGccATGATGACCTACGAATGG–3′ | TAGCopenhagen, used for generation of Glu466Ala variant |
| Oligo14_Glu466Ala_Rev | 5′–TTCGTAGGTCATCATggCTTGGTCCACTAAAGAAATGTC–3′ | TAGCopenhagen, used for generation of Glu466Ala variant |
| Oligo15_Asp52Glu_Fwd | 5′–GTGCCCGGTGGAGAgTCCGGAGAGCTGATTAC–3′ | TAGCopenhagen, used for generation of Asp52Glu variant |
| Oligo16_Asp52Glu_Rev | 5′–GTAATCAGCTCTCCGGAcTCTCCACCGGGCAC–3′ | TAGCopenhagen, used for generation of Asp52Glu variant |
| Oligo17_Glu582Gln_Fwd | 5′– | TAGCopenhagen, used for |

| | | |
|---|---|---|
| | GACCCCTCCAGCTGGcAGTCCGTGGCC–3′ | generation of Glu582Gln variant |
| Oligo18_Glu582Gln_Rev | 5′–ATTGGCCACGGACTgCCAGCTGGAGGG–3′ | TAGCopenhagen, used for generation of Glu582Gln variant |
| Oligo19_Val494Ala_Fwd | 5′–GCAATAGGTGGAACCCCGccGAGGGCATTCTGCCCAGC–3′ | TAGCopenhagen, used for generation of Val494Ala variant |
| Oligo20_Val494Ala_Rev | 5′–GCTGGGCAGAATGCCCTCggCGGGGTTCCACCTATTGC–3′ | TAGCopenhagen, used for generation of Val494Ala variant |
| Oligo21_R21A-R22A_Fwd | 5′–GCTGTGAGGGTGGGTTTAgccgccAGCGGCGGAATTAGGGG–3′ | TAGCopenhagen, used for generation of R21A/R22A double mutant |
| Oligo22_R21A-R22A_Rev | 5′–CCCCTAATTCCGCCGCTggcggcTAAACCCACCCTCACAGC–3′ | TAGCopenhagen, used for generation of R21A/R22A double mutant |
| Oligo23_R21A-R22S_Fwd | 5′–GCTGTGAGGGTGGGTTTAgccagcAGCGGCGGAATTAGGGG–3′ | TAGCopenhagen, used for generation of R21A/R22S double mutant |
| Oligo24_R21A-R22S_Rev | 5′–CCCTAATTCCGCCGCTgctggcTAAACCCACCCTCACAGC–3′ | TAGCopenhagen, used for generation of R21A/R22S double mutant |
| Oligo25_Cys544Ser_Fwd | 5′–GGCCTTGGGGCTCTagcGATAAGCTGGTGCC–3′ | TAGCopenhagen, used for generation of Cys544Ser variant |
| Oligo26_Cys544Ser_Rev | 5′–GGCACCAGCTTATCgctAGAGCCCCAAGGCC–3′ | TAGCopenhagen, used for generation of Cys544Ser variant |
| Oligo27_Cys522Ser_Fwd | 5′–GGAGACCTTCGTGagcATCGGCATCCACG–3′ | TAGCopenhagen, used for generation of Cys522Ser variant |
| Oligo28_Cys522Ser_Rev | 5′–GTGGATGCCGATgctCACGAAGGTCT | TAGCopenhagen, used for generation of Cys522Ser |

| | | |
|---|---|---|
| | CC–3′ | variant |
| Oligo29_Arg134Ala_Fwd | 5′–CCGGAGTGTTTAGCTTTTCTgccCTGACATGGCAGTGGTCCATCG–3′ | TAGCopenhagen, used for generation of Arg134Ala variant |
| Oligo30_Arg134Ala_Rev | 5′–CGATGGACCACTGCCATGTCAGggcAGAAAAGCTAAACACTCCGG–3′ | TAGCopenhagen, used for generation of Arg134Ala variant |
| Oligo31_His526Ala_Fwd | 5′–CGTGTGCATCGGCATCgccGAGGGCGATCCTAC–3′ | TAGCopenhagen, used for generation of His526Ala variant |
| Oligo32_His526Ala_Rev | 5′–GTAGGATCGCCCTCggcGATGCCGATGCACACG–3′ | TAGCopenhagen, used for generation of His526Ala variant |
| Oligo33_Glu582Ala_Fwd | 5′–CCCTCCAGCTGGgccTCCGTGGCCAATGAGG–3′ | TAGCopenhagen, used for generation of Glu582Ala variant |
| Oligo34_Glu582Ala_Rev | 5′–CCTCATTGGCCACGGAggcCCAGCTGGAGGG–3′ | TAGCopenhagen, used for generation of Glu582Ala variant |
| Oligo35_VectorAsnLadder_Fwd | 5′–GCACCTCAGAAAGGAGCTGC–3′ | TAGCopenhagen, used for vector amplification for asparagine ladder variant |
| Oligo36_VectorAsnLadder_Rev | 5′–ACGAGCTTGCCACATCTCC–3′ | TAGCopenhagen, used for vector amplification for asparagine ladder variant |

**Table S4. Cryo-EM data collection, refinement and validation statistics.**

| | **Binary complex** TMEM260 with Far-P-Man<br><br>EMD-54178)<br>(PDB 9RQL) | **Binary complex** TMEM260 with DoI-P-Man<br><br>(EMD-54180)<br>(PDB 9RQN) | **Ternary complex** TMEM260 with Dol-P-Man and PLNXB2-IPT1 peptide<br>(EMD-54179)<br>(PDB 9RQM) |
|---|---|---|---|
| **Data collection and processing** | | | |
| Magnification | 105000 | 105000 | 105000 |
| Voltage (kV) | 300 | 300 | 300 |
| Electron exposure (e–/Å$^2$) | 60.1 | 60.5 | 49.8 |
| Defocus range (μm) | 0.5-2.0 | 0.5-2.0 | 0.5-2.0 |
| Pixel size (Å) | 0.8238 | 0.8238 | 0.8238 |
| Movies collected | 12964 | 12967 | 21886 |
| Initial particle images (no.) | 3130683 | 2290548 | 3296713 |
| Final particle images (no.) | 332040 | 635355 | 271859 |
| Map resolution (Å)<br>FSC threshold | 2.65<br>0.143 | 2.94<br>0.143 | 3.06<br>0.143 |
| Map resolution range (Å) | 1.78-8.90 | 2.52-8.95 | 2.66-8.10 |
| **Refinement** | | | |
| Model resolution (Å)<br>FSC threshold | 2.6<br>0.143 | 2.9<br>0.143 | 3.0<br>0.143 |
| Model resolution range (Å) | 1.78-8.90 | 2.52-8.95 | 2.66-8.10 |
| Map sharpening *B* factor (Å$^2$) | -103.6 | -118.0 | -118.9 |
| Model composition<br>Non-hydrogen atoms<br>Protein residues<br>Ligands | <br>5320<br>676<br>2 | <br>5331<br>708<br>2 | <br>5332<br>676<br>2 |
| *B* factors (Å$^2$)<br>Protein<br>Ligand | <br>106.58<br>115.86 | <br>58.60<br>63.57 | <br>98.76<br>119.69 |
| R.M.S. deviations<br>Bond lengths (Å)<br>Bond angles (°) | <br>0.008<br>0.855 | <br>0.006<br>0.890 | <br>0.007<br>0.904 |
| Validation<br>MolProbity score<br>Clashscore<br>Poor rotamers (%) | <br>1.18<br>3.99<br>0.18 | <br>1.06<br>2.09<br>0.18 | <br>1.38<br>3.75<br>0.35 |
| Ramachandran plot<br>Favored (%)<br>Allowed (%)<br>Disallowed (%) | <br>98.36<br>1.49<br>0.15 | <br>97.62<br>2.23<br>0.15 | <br>96.59<br>3.27<br>0.14 |

### 2. Supplementary Figures

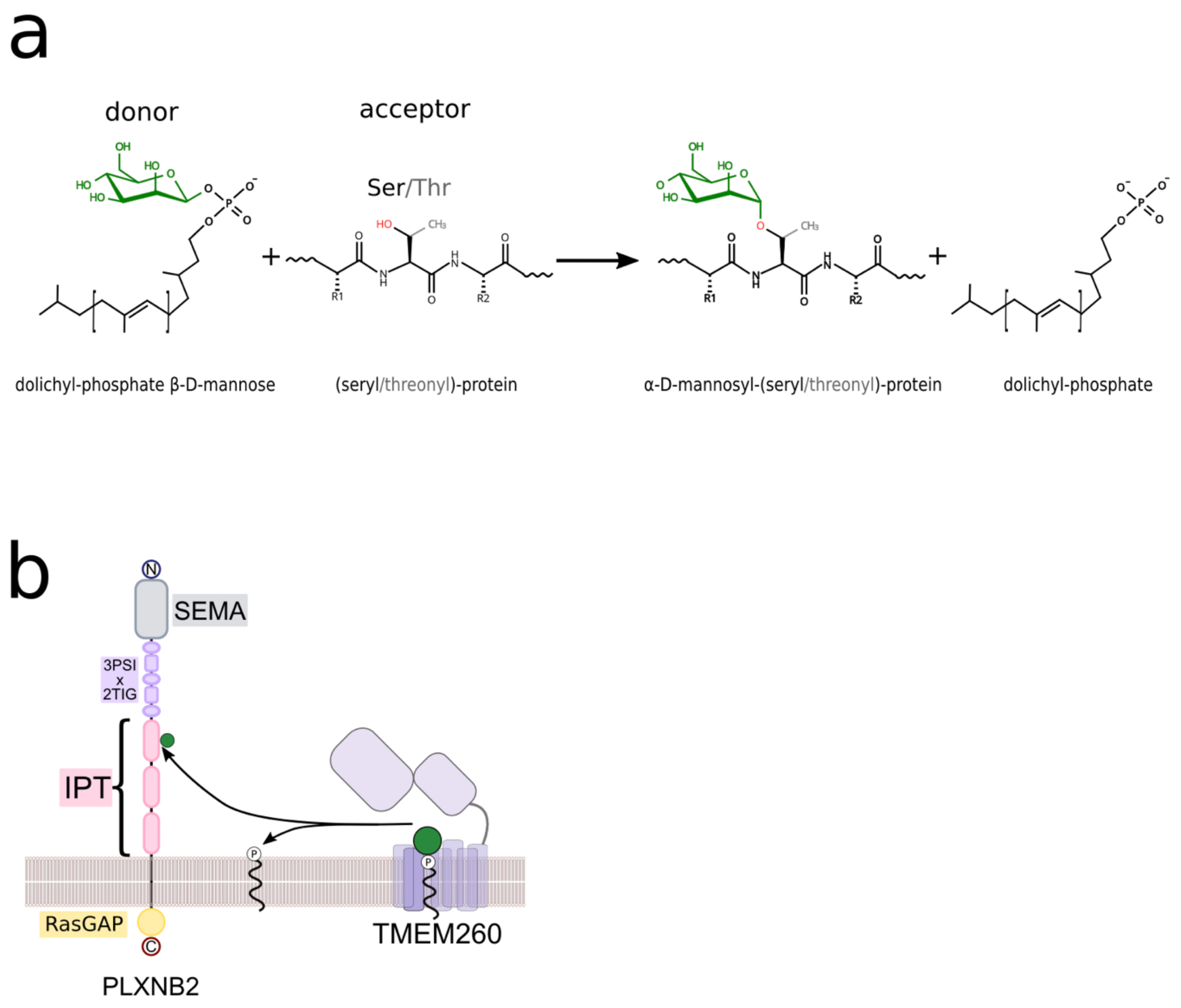

**Fig. S1. O-mannosyltransferase reaction and TMEM260 substrate specificity. a,** O-mannosyltransferases catalyze the transfer of α-mannose from dolichyl-β-D-mannosyl phosphate to serine or threonine hydroxyl groups in protein acceptors. **b,** Schematic of TMEM260-mediated O-mannosylation of PLXNB2 IPT domains.

a

MSPHGDGRGQAQGRAVRVGLRRSGGIRGGVAVFAAVAAVFTFTLPPSVPGGDSGELITAAHELGVAHPPGYPLFTLVAKLAITLFPFGSIAYRVNLLCGL (100)
FGAVAASLLFFTVFRLSGSSAGGILAAGVFSFSRLTWQWSIAAEVFSLNNLFVGLLMALTVHFEEAATAKERSKVAKIGAFCCGLSLCNQHTIILYVLCI (200)
IPWILFQLLKKKELSLGSLLKLSLYFSAGLLPYVHLPISSYLNHARWTWGDQTTLQGFLTHFLREEYGTFSLAKSEIGSSMSEILLSQVTNMRTELSFNI (300)
QALAVCANICLATKDRQNPSLVWLFTGMFCIYSLFFAWRANLDISKPLFMGVVERFWMQSNAVVAVLAGIGLAAVVSETNRVLNSNGLQCLEWLSATLFV (400)
VYQIYSNYSVCDQRTNYVIDKFAKNLLTSMPHDAIILLRGDLPGNSLRYMHYCEGLRPDISLVDQEMMTYEWYLPKMAKHLPGVNFPGNRWNPVEGILPS (500)
GMVTFNLYHFLEVNKQKETFVCIGIHEGDPTWKKNYSLWPWGSCDKLVPLEIVFNPEEWIKLTKSIYNWTEEYGRFDPSSWESVANEEMWQARMKTPFFI (600)
FNLAETAHMPSKVKAQLYAQAYDLYKEIVYLQKEHPVNWHKNYAIACERMLRLQARDADPEVLLSETIRHFRLYSQKAPNDPQQADILGALKHLRKELQS (700)
LRNRKNV(707)DYKDHDGDYKDHDIDYKDDDDK

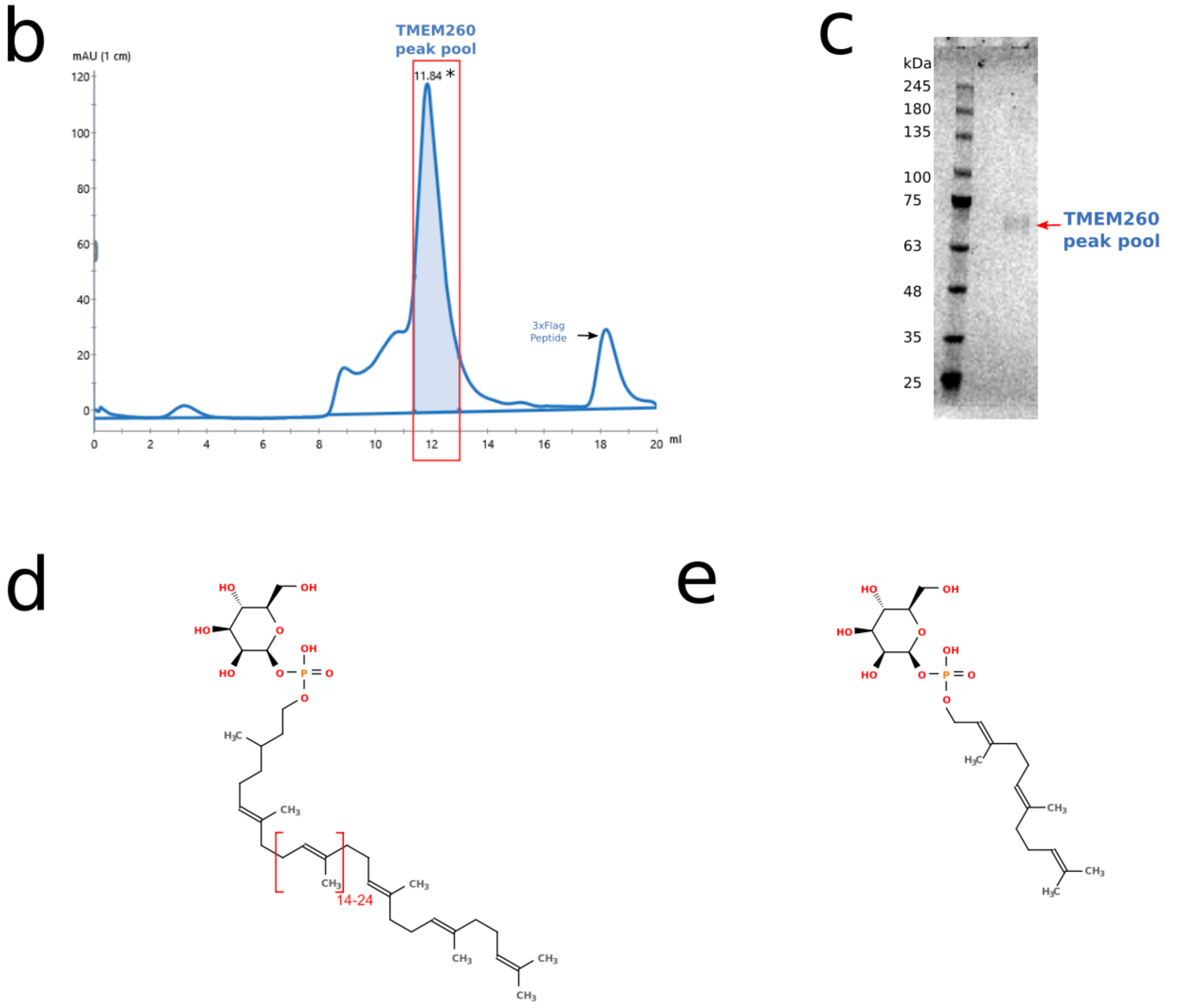

β-D-mannosyl dolichyl phosphate
(Dol-P-Man)

β-D-mannosyl farnesyl phosphate
(Far-P-Man)

**Fig. S2. TMEM260-3×FLAG construct, purification and donor structures. a,** Amino-acid sequence of full-length TMEM260 (black) with the C-terminal 3×FLAG tag (red). **b,** Size-exclusion chromatogram (Superdex 200 Increase 10/300) of purified TMEM260-3×FLAG expressed in Expi293 cells used for cryo-EM. **c,** Coomassie-stained SDS-PAGE of pooled peak fractions. **d-e,** Chemical structures of Dol-P-Man and the donor analogue Far-P-Man.

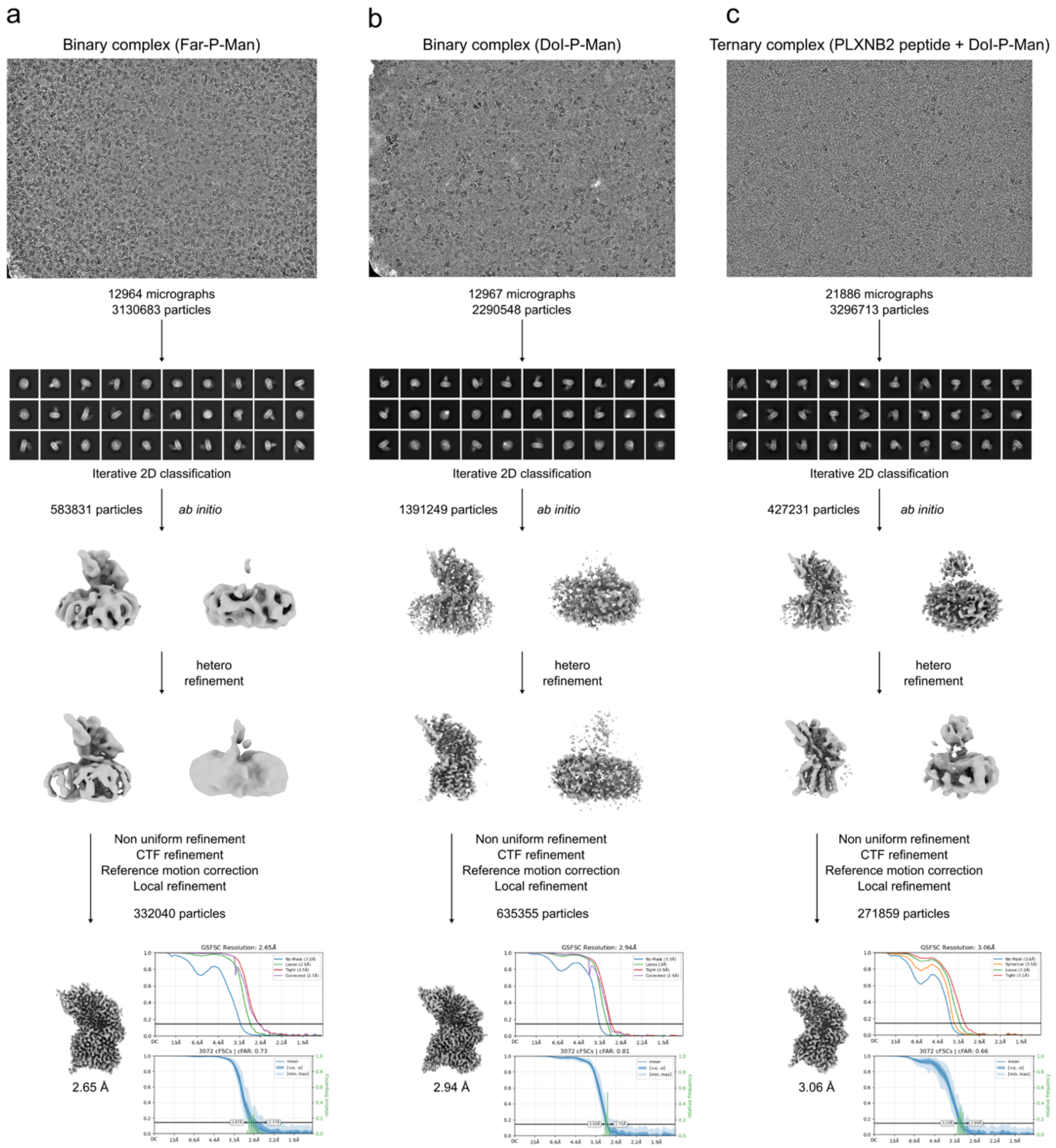

**Fig. S3. Cryo-EM data processing.** Cryo-EM data processing workflows for (**a**) TMEM260:Far-P-Man, (**b**) TMEM260:Dol-P-Man, and (**c**) TMEM260:Dol-P-Man:PLXB2-IPT1. All datasets were processed using cryoSPARC. Image processing began with particle picking, followed by iterative 2D classification, ab initio reconstruction, and heterogeneous refinement. This was followed by non-uniform refinement, CTF refinement, motion correction, and local refinement. Gold-standard FSC and cFSC analyses are shown for all three complexes.

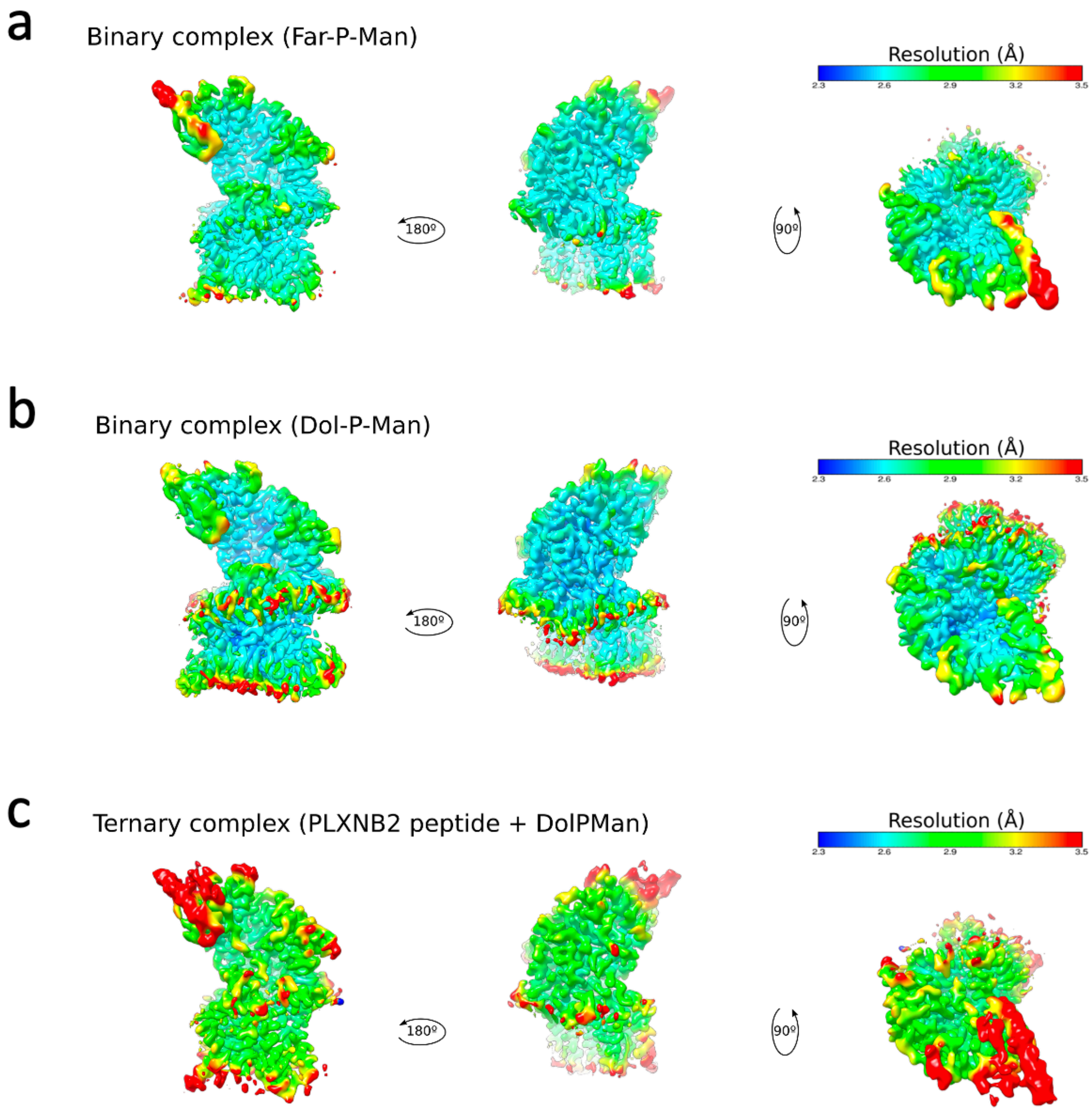

**Fig. S4. Local map resolution.** Three orthogonal views of the cryo-EM maps of (**a**) TMEM260:Far-P-Man binary complex, (**b**) TMEM260:Dol-P-Man binary complex and (**c**) TMEM260:Dol-P-Man:PLXB2-IPT1 ternary complex colored as a function of resolution.

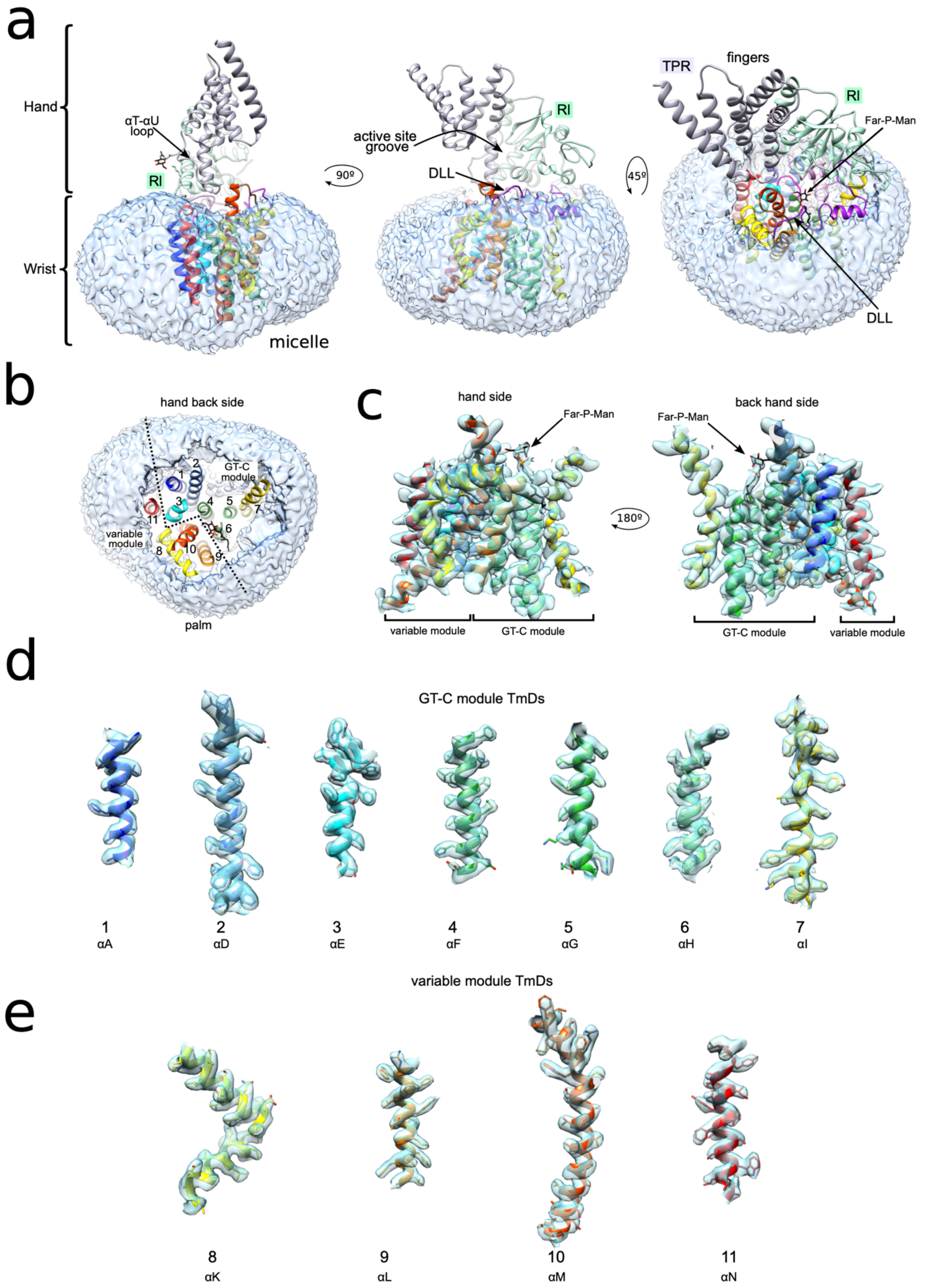

**Fig. S5. The transmembrane region of the TMEM260:Far-P-Man binary complex. a,** Three views of the cryo-EM reconstruction of TMEM260 bound to Far-P-Man, displayed within the detergent micelle difference map. The micelle boundary delineates the transmembrane (wrist) and

ER-luminal (hand) regions. **b,** Top view of the micelle with the ER-luminal lobe removed, revealing the relative arrangement of the transmembrane helices (numbered) with the GT-C and variable modules (separated by a discontinuous trace). The donor analogue binds at the interface between both modules and its isoprenoid chain extends into the micelle between TMHs 6 and 9, delineating a putative donor-entry trajectory toward the active site. **c,** Overviews of the cryo-EM density and corresponding atomic model for the transmembrane region. **d-e,** Local density views for individual transmembrane helices, denoting map-model agreement and side-chain resolvability for unambiguous assignment.

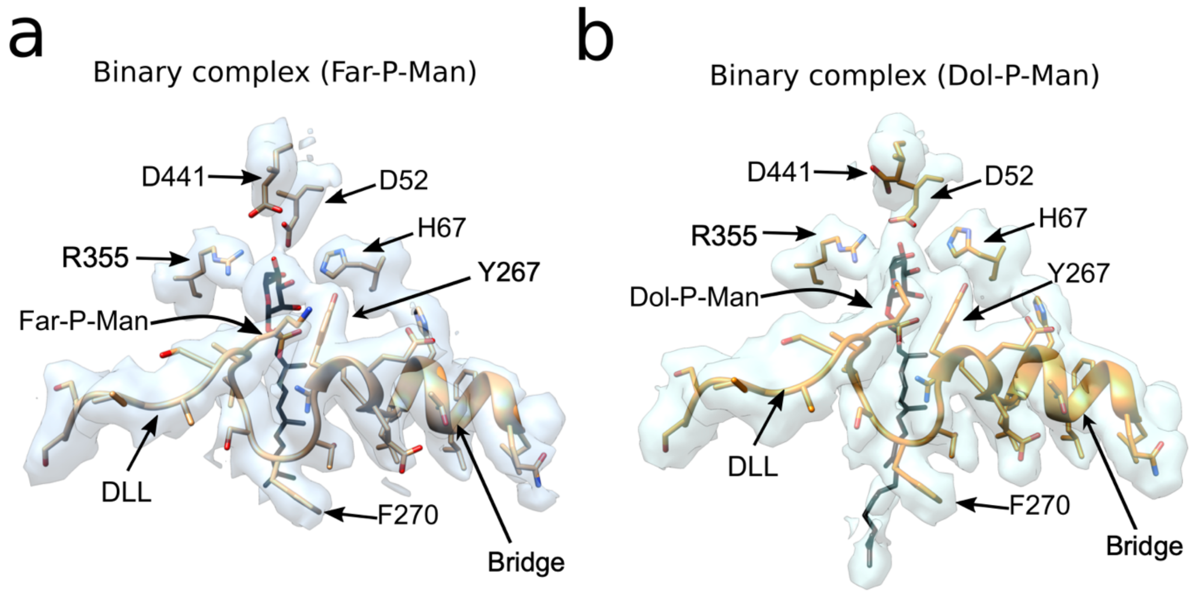

**Fig. S6. Active site architecture in TMEM260 binary complexes. a,** Close-up of the donor-binding pocket of TMEM260 bound to the donor analogue Far-P-Man (2.7 Å resolution). **b,** Close-up of the donor-binding pocket of TMEM260 bound to the native donor Dol-P-Man (2.9 Å resolution). Side-by-side comparison reveals that the active site adopts an essentially identical conformation in both binary complexes. The donor-loading loop (DLL) and the bridge helix display closely matched density in the two reconstructions. In both cases, D52 contacts and helps orient the sugar moiety, whereas D441 remains disengaged.

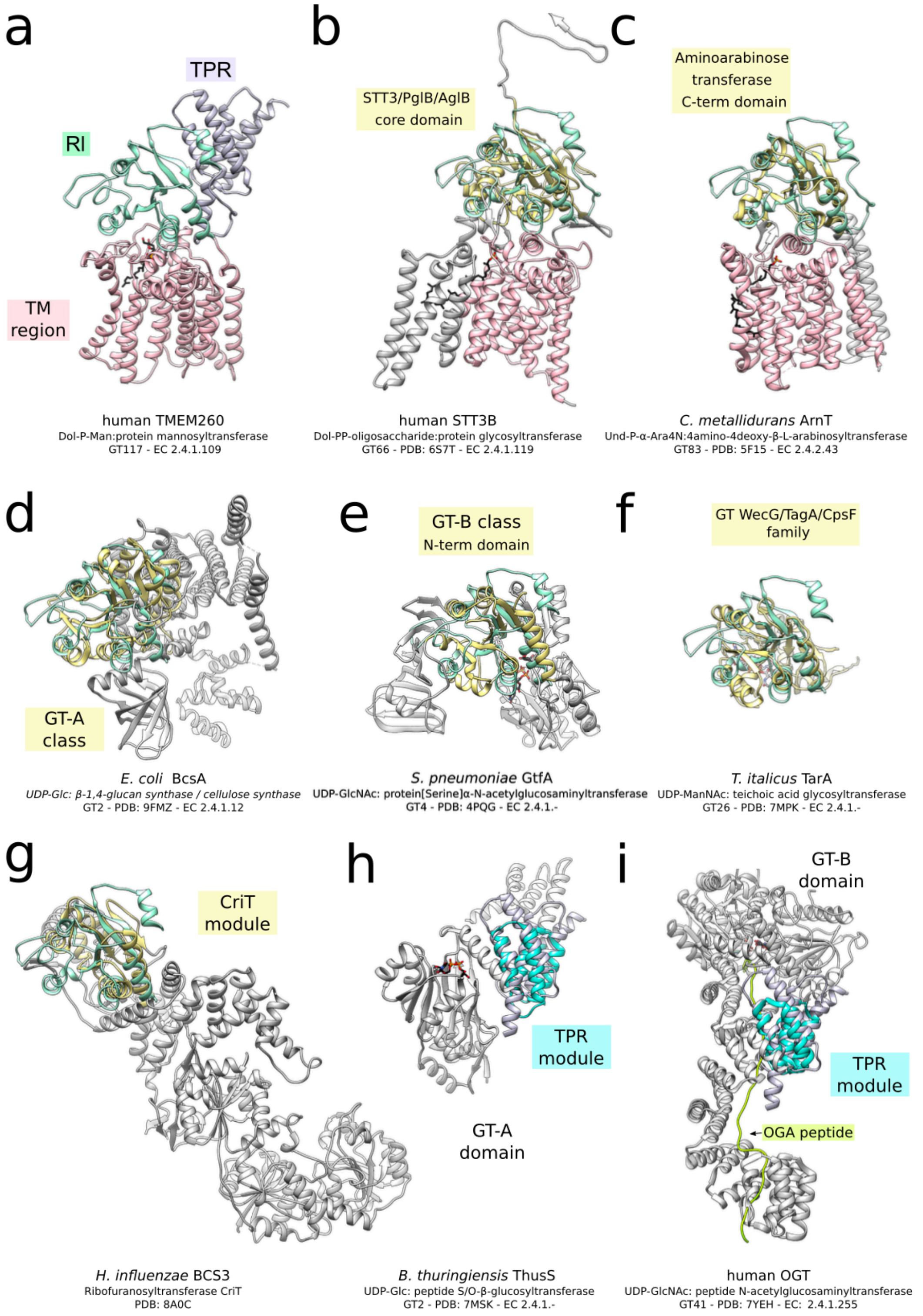

**Fig. S7. Structural comparisons of TMEM260 with representative glycosyltransferases. a,** Reference view of TMEM260 (back of the active site), with the Rl domain in green, the TPR domain in lavender, and the transmembrane region pink. Superpositions of isolated TMEM260 luminal domains (oriented as in **a**) with structural matches identified by DALI. TMEM260 regions aligned to matched domains are shown in pale yellow (Rl) or cyan (TPR); non-superposed regions are shown in grey. Enzyme name, function, CAZy family and EC classification are indicated in each panel. **b-c,** GT-C enzymes displaying distant similarity to the TMEM260 Rl domain; homologous transmembrane regions are shown in pink (note the alternate orientation relative to

**a**). **d-e,** Superpositions of the Rl domain with canonical Leloir-type GT-A and GT-B folds. **f-g,** Comparisons with TarA and the BCS3 ribofuranosyltransferase module CriT, which lack formal CaZy structural classification but exhibit distant similarity, consistent with convergent evolution. **h-i,** Matches involving the TMEM260 TPR domain and peptide-glycosylating Leloir-type enzymes, including O-GlcNAc transferase (OGT) in **i**, shown bound to an O-GlcNAcase (OGA) peptide.

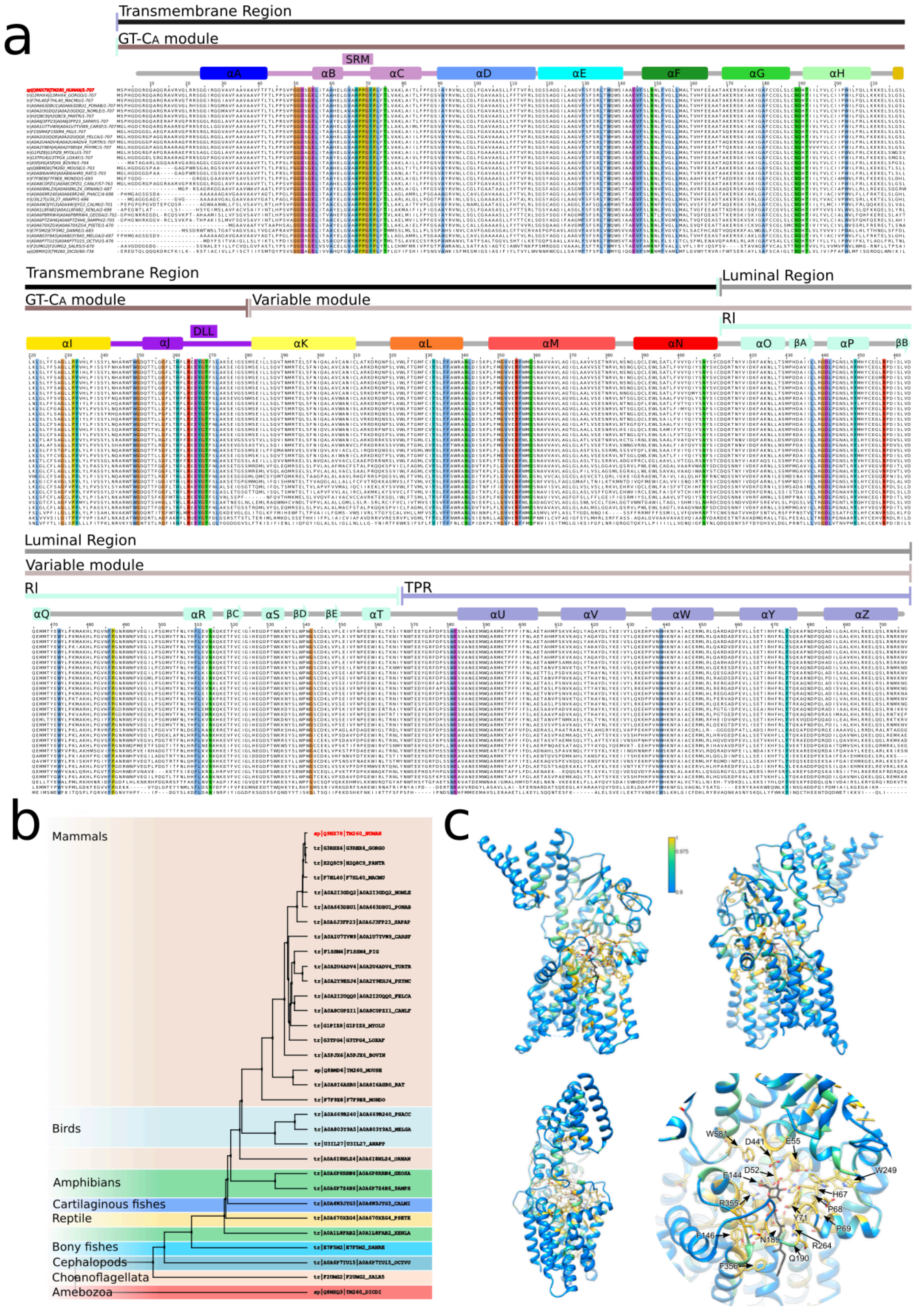

**Fig. S8. Conservation analysis of TMEM260 across species. a,** Sequence alignment of human TMEM260 with selected homologues with high-confidence AlphaFold models. Regions corresponding to gaps in the human sequence were removed for clarity. Strictly conserved residues (Clustal coloring) cluster predominantly within the transmembrane region, with reduced conservation toward the TPR domain, consistent with preservation of catalytic function and species-specific variation in acceptor binding. **b,** Neighbor-joining phylogenetic tree of the aligned sequences. **c,** Views of TMEM260:Far-P-Man complex, including an active-site close-up, with strictly conserved residues in yellow and highly conserved residues in green (threshold ≥ 0.975).

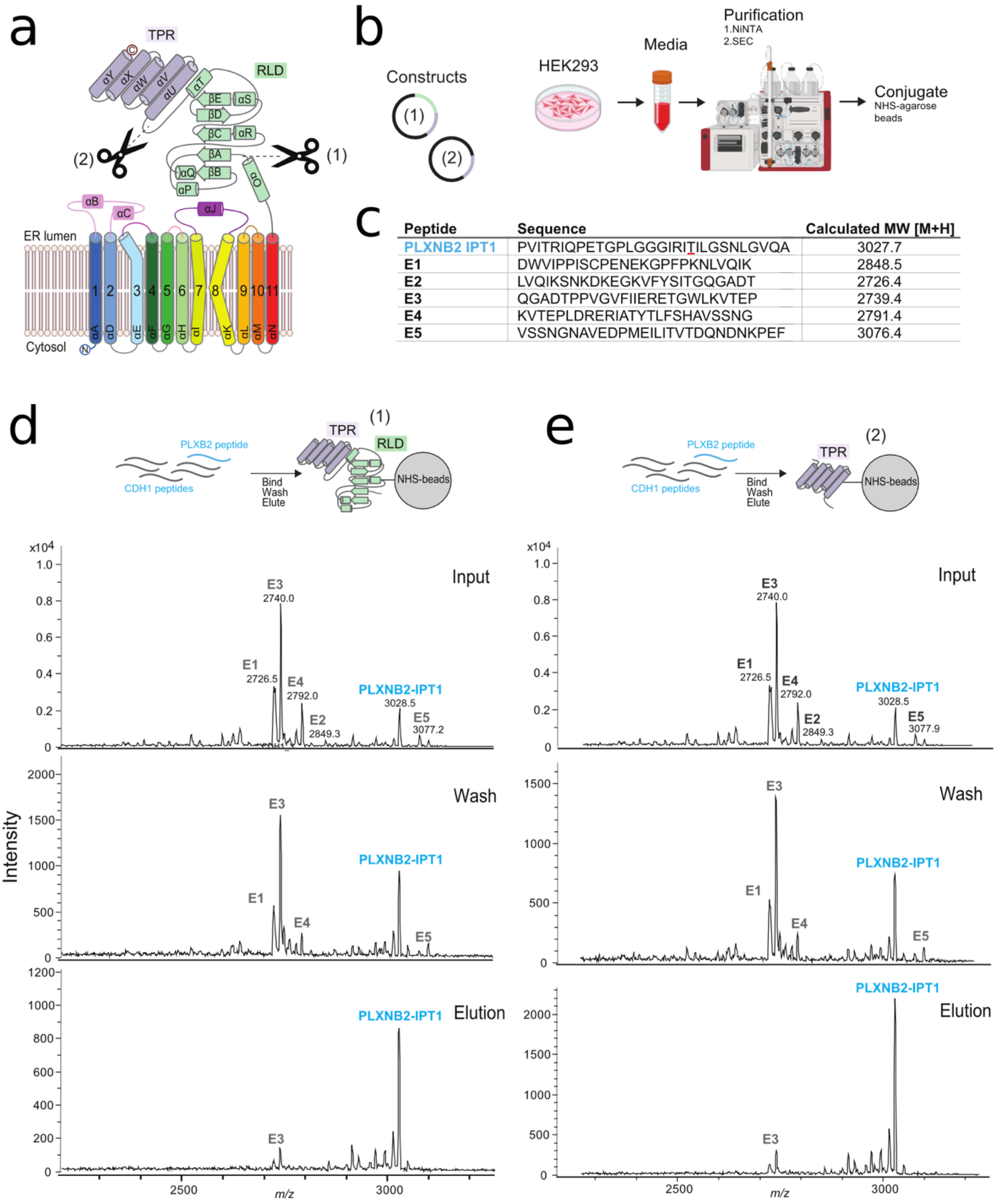

**Fig. S9. PLXNB2-IPT1 pull-down by the ER-luminal domains of TMEM260. a,** Topology diagram of TMEM260 highlighting the ER-luminal Rl-TPR (construct 1) and TPR-only (construct 2) truncations used in the pull-down assays. **b,** Schematic of the production and agarose conjugation of the two constructs, expressed as superfolder GFP (sfGFP)-mCherry fusion proteins in HEK293 cells. **c,** Amino acid sequences of the PLXNB2-IPT1 peptide and control 25-mer peptides derived from human E-cadherin, with calculated molecular weights (MW). **d-e,** MALDI-TOF analysis of peptide pulldowns using beads functionalized with the TMEM260 luminal truncations (Rl-TPR or TPR-only). The TPR region is necessary and sufficient for recruitment of the PLXNB2-IPT1 peptide.

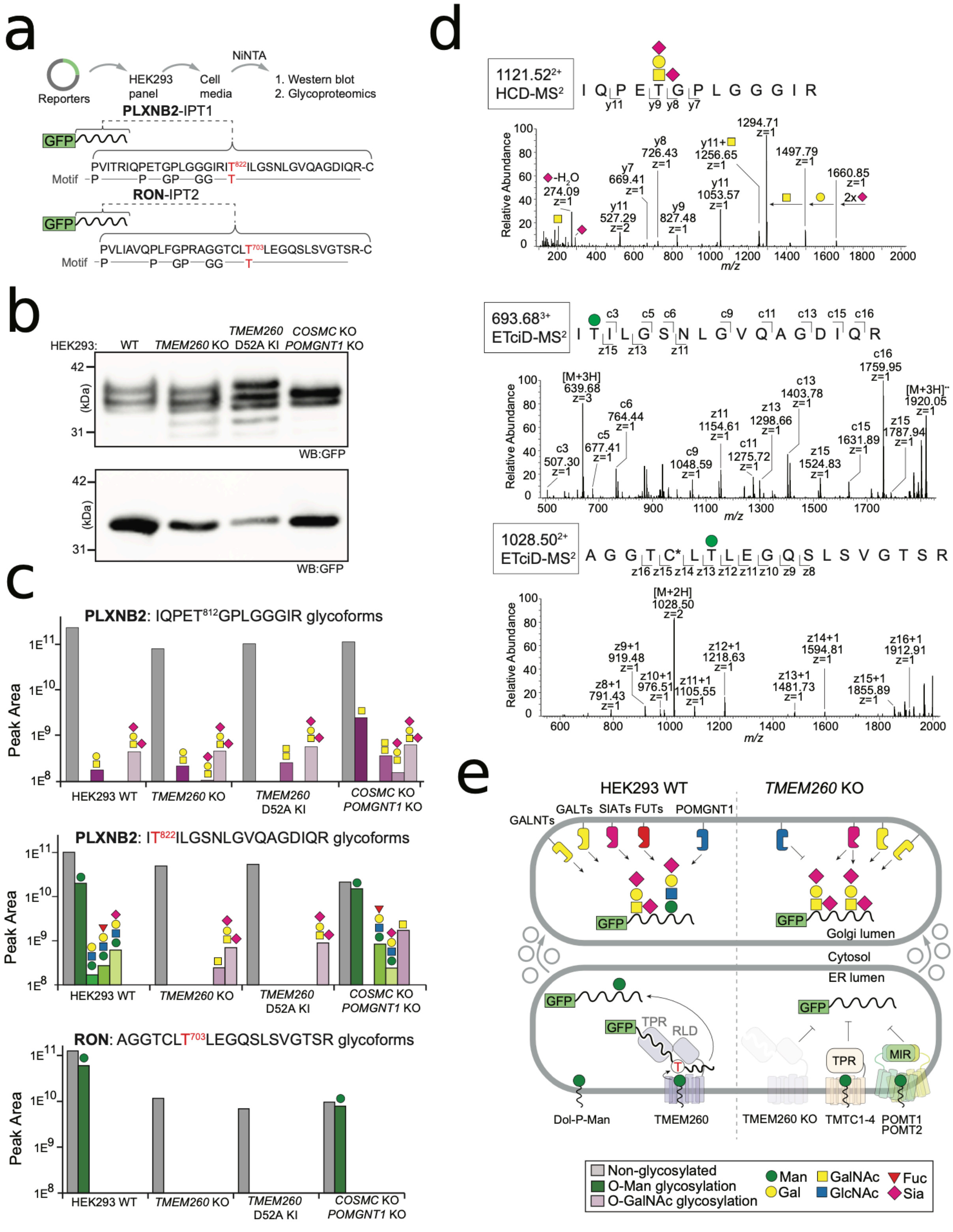

**Fig. S10. TMEM260-specific O-mannosylation of Plexin-B2 and RON receptor peptides fused to GFP-reporters. a,** Design and strategy for the in cellulo assay using sfGFP-fused reporters bearing a N-terminal ER signal peptide and 12xHis tag (not shown) for Ni-NTA purification. Reporters contain either a 35-mer peptide from PLXB2-IPT1 domain encompassing the T822 glycosylation site (red) or a 32-mer peptide from the RON-IPT2 domain encompassing the T702 glycosylation site (red). The TMEM260 TPR-recognition motif

(PXnPXnGPXnGGXnT) is indicated below each sequence. Reporters were transiently expressed in wild-type (WT) HEK293 cells and in glycoengineered HEK293 cells lacking TMEM260 (KO) or expressing a catalytically inactive TMEM260 KO/TMEM260 D52A knock-in (KI), followed by Ni-NTA purification, immunoblotting and bottom-up glycoproteomics. COSMC/POMGNT1 KO "SimpleCells" served as a glycoengineering control. **b,** Anti-GFP immunoblots reveal multiple glycoforms for the sfGFP-PLXNB2-IPT1 reporter (top), whereas the sfGFP-RON-IPT2 reporter migrates as a single predominant species (bottom). The calculated molecular mass of unglycosylated reporters is ~36 kDa. **c,** Label-free quantification of glycopeptide abundances based on area-under-the-curve (AUC) measurements for MS/MS-identified glycoforms. TMEM260-dependent sites (T822 in PLXNB2 and T703 in RON) are highlighted in red. Tryptic digests of Ni-NTA-purified reporters from conditioned media were analyzed by data-dependent acquisition (DDA) and bottom-up glycoproteomics employing Collision-Induced Dissociation (CID), Higher-Energy Collision Dissociation (HCD), and hybrid Electron-Transfer Collision-Induced Dissociation (ETciD) fragmentation modes on an Orbitrap Fusion Lumos instrument. Bar charts show peptide-level abundances (y-axis; peak areas) grouped by cellular background. Glycans are depicted using Society for Glycobiology (SFG) standardized nomenclature of colored geometric shapes: green circle for mannose, yellow circle for galactose, blue square for N-acetylglucosamine, yellow square for N-acetylgalactosamine, red triangle for fucose and pink diamond for sialic acid (NeuAc). Abundance for non-glycosylated peptides is illustrated with grey bars, O-Man glycoforms (green shades) and O-GalNAc glycoforms (purple shades). **d,** Representative, partially annotated MS/MS spectra used to identify peptide backbones and glycan moieties. ETciD spectra unambiguously localize O-mannose to T822 (PLXNB2) and T703 (RON). **e,** Model summarizing the experimental findings. In WT cells (left), the sfGFP reporter enters the ER lumen, where the extended C-terminal peptide is recruited by TMEM260 and the target threonine is positioned for O-mannosylation. Following ER exit, the O-Man monosaccharide is partially elaborated in the Golgi into core-M1 glycans by POMGNT1, GALT, FUT and SIAT enzymes; in parallel, O-GalNAc glycosylation by GALNTs occurs at T812 of the sfGFP-PLXNB2-IPT1 reporter and is further extended into sialyl-T and disialyl-T structures before secretion. In TMEM260-deficient cells (right), reporters are not O-mannosylated despite the presence of TMTC1-4 and POMT1/2, demonstrating strict substrate specificity. Instead, both T812 and T822 remain available for Golgi-initiated O-GalNAc glycosylation by GALNTs and subsequent elaboration into disialyl-T glycans by GALT and SIAT enzymes.

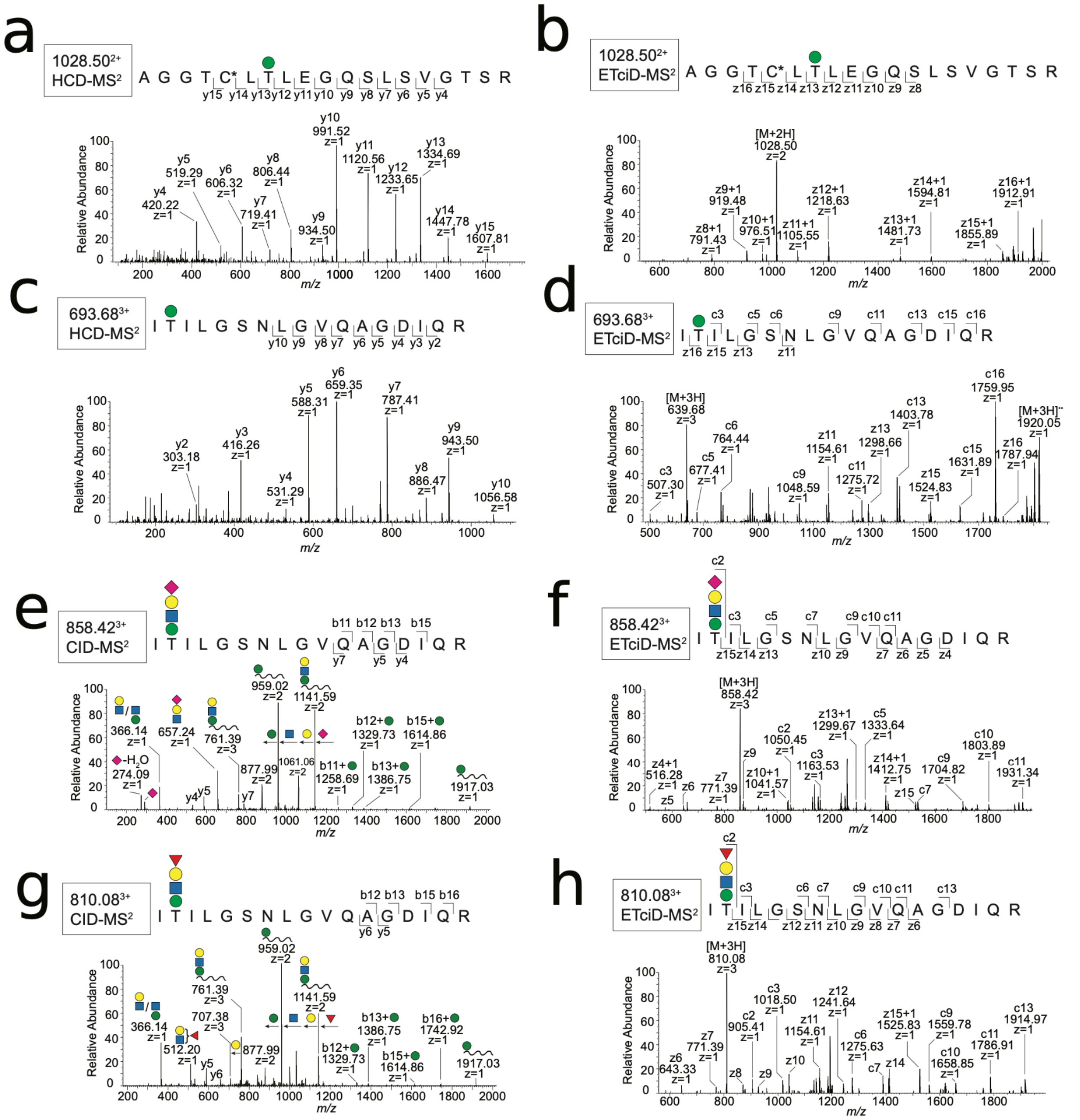

a
1028.50²⁺ HCD-MS²
A G G T C* L T L E G Q S L S V G T S R
y15 y14 y13 y12 y11 y10 y9 y8 y7 y6 y5 y4
y4 420.22 z=1
y5 519.29 z=1
y6 606.32 z=1
y7 719.41 z=1
y8 806.44 z=1
y9 934.50 z=1
y10 991.52 z=1
y11 1120.56 z=1
y12 1233.65 z=1
y13 1334.69 z=1
y14 1447.78 z=1
y15 1607.81 z=1
Relative Abundance
m/z
b
1028.50²⁺ ETciD-MS²
A G G T C* L T L E G Q S L S V G T S R
z16 z15 z14 z13 z12 z11 z10 z9 z8
[M+2H] 1028.50 z=2
z8+1 791.43 z=1
z9+1 919.48 z=1
z10+1 976.51 z=1
z11+1 1105.55 z=1
z12+1 1218.63 z=1
z13+1 1481.73 z=1
z14+1 1594.81 z=1
z15+1 1855.89 z=1
z16+1 1912.91 z=1
Relative Abundance
m/z
c
693.68³⁺ HCD-MS²
I T I L G S N L G V Q A G D I Q R
y10 y9 y8 y7 y6 y5 y4 y3 y2
y2 303.18 z=1
y3 416.26 z=1
y4 531.29 z=1
y5 588.31 z=1
y6 659.35 z=1
y7 787.41 z=1
y8 886.47 z=1
y9 943.50 z=1
y10 1056.58 z=1
Relative Abundance
m/z
d
693.68³⁺ ETciD-MS²
I T I L G S N L G V Q A G D I Q R
c3 c5 c6 c9 c11 c13 c15 c16
z16 z15 z13 z11
[M+3H] 639.68 z=3
c3 507.30 z=1
c5 677.41 z=1
c6 764.44 z=1
c9 1048.59 z=1
z11 1154.61 z=1
c11 1275.72 z=1
z13 1298.66 z=1
c13 1403.78 z=1
z15 1524.83 z=1
c15 1631.89 z=1
c16 1759.95 z=1
z16 1787.94 z=1
[M+3H]⁺⁺ 1920.05 z=1
Relative Abundance
m/z
e
858.42³⁺ CID-MS²
I T I L G S N L G V Q A G D I Q R
b11 b12 b13 b15
y7 y5 y4
-H₂O 274.09 z=1
366.14 z=1
657.24 z=1
761.39 z=3
877.99 z=2
959.02 z=2
1061.06 z=2
1141.59 z=2
b11+ 1258.69 z=1
b12+ 1329.73 z=1
b13+ 1386.75 z=1
b15+ 1614.86 z=1
1917.03 z=1
y4 y5 y7
Relative Abundance
m/z
f
858.42³⁺ ETciD-MS²
I T I L G S N L G V Q A G D I Q R
c2 c3 c5 c7 c9 c10 c11
z15 z14 z13 z10 z9 z7 z6 z5 z4
[M+3H] 858.42 z=3
z4+1 516.28 z=1
z5
z6
z7 771.39 z=1
z9
c2 1050.45 z=1
z10+1 1041.57 z=1
c3 1163.53 z=1
z13+1 1299.67 z=1
c5 1333.64 z=1
z14+1 1412.75 z=1
z15
c7
c9 1704.82 z=1
c10 1803.89 z=1
c11 1931.34 z=1
Relative Abundance
m/z
g
810.08³⁺ CID-MS²
I T I L G S N L G V Q A G D I Q R
b12 b13 b15 b16
y6 y5
366.14 z=1
512.20 z=1
y5
y6
707.38 z=3
761.39 z=3
877.99 z=2
959.02 z=2
1141.59 z=2
b12+ 1329.73 z=1
b13+ 1386.75 z=1
b15+ 1614.86 z=1
b16+ 1742.92 z=1
1917.03 z=1
Relative Abundance
m/z
h
810.08³⁺ ETciD-MS²
I T I L G S N L G V Q A G D I Q R
c2 c3 c6 c7 c9 c10 c11 c13
z15 z14 z12 z11 z10 z9 z8 z7 z6
[M+3H] 810.08 z=3
z6 643.33 z=1
z7 771.39 z=1
z8
z9
c2 905.41 z=1
c3 1018.50 z=1
z10
z11 1154.61 z=1
z12 1241.64 z=1
c6 1275.63 z=1
c7
z14
z15+1 1525.83 z=1
c9 1559.78 z=1
c10 1658.85 z=1
c11 1786.91 z=1
c13 1914.97 z=1
Relative Abundance
m/z

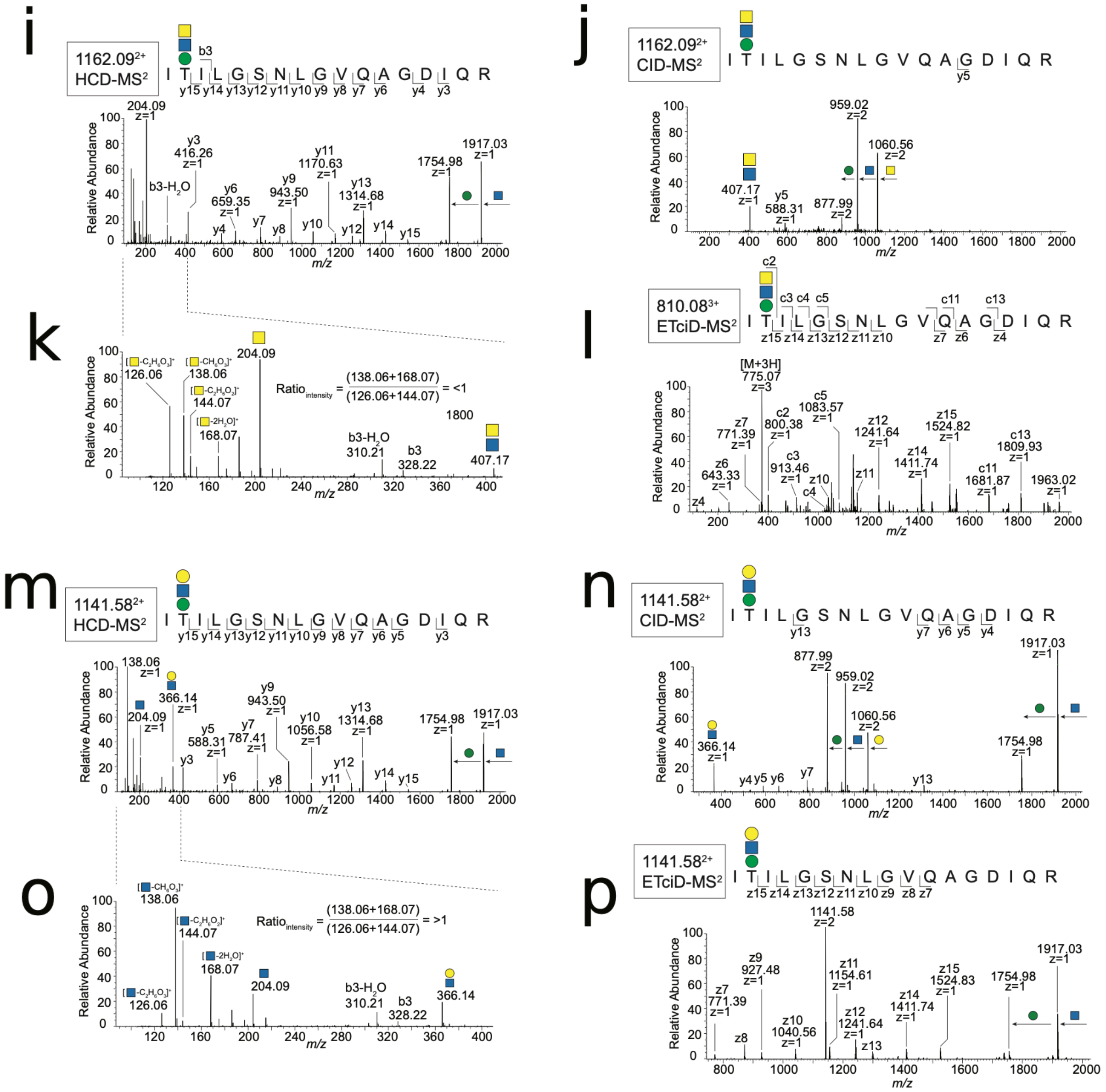
i
1162.09²⁺
HCD-MS²
I T I L G S N L G V Q A G D I Q R
b3
y15 y14 y13 y12 y11 y10 y9 y8 y7 y6 y4 y3
204.09 z=1
y3 416.26 z=1
b3-H2O
y6 659.35 z=1
y9 943.50 z=1
y11 1170.63 z=1
y13 1314.68 z=1
1754.98 z=1
1917.03 z=1
Relative Abundance
m/z
j
1162.09²⁺
CID-MS²
I T I L G S N L G V Q A G D I Q R
y5
959.02 z=2
1060.56 z=2
407.17 z=1
y5 588.31 z=1
877.99 z=2
k
[-C2H4O2]⁺ 126.06
[-CH6O3]⁺ 138.06
[-C2H4O2]⁺ 144.07
[-2H2O]⁺ 168.07
204.09
Ratio intensity = (138.06+168.07)/(126.06+144.07) = <1
b3-H2O 310.21
b3 328.22
407.17
l
810.08³⁺
ETciD-MS²
I T I L G S N L G V Q A G D I Q R
c2 c3 c4 c5 c11 c13
z15 z14 z13 z12 z11 z10 z7 z6 z4
[M+3H] 775.07 z=3
z7 771.39 z=1
c2 800.38 z=1
c5 1083.57 z=1
z12 1241.64 z=1
z15 1524.82 z=1
c13 1809.93 z=1
z6 643.33 z=1
c3 913.46 z=1
z14 1411.74 z=1
c11 1681.87 z=1
1963.02 z=1
m
1141.58²⁺
HCD-MS²
I T I L G S N L G V Q A G D I Q R
y15 y14 y13 y12 y11 y10 y9 y8 y7 y6 y5 y3
138.06 z=1
366.14 z=1
204.09 z=1
y9 943.50 z=1
y5 588.31 z=1
y7 787.41 z=1
y10 1056.58 z=1
y13 1314.68 z=1
1754.98 z=1
1917.03 z=1
n
1141.58²⁺
CID-MS²
I T I L G S N L G V Q A G D I Q R
y13 y7 y6 y5 y4
877.99 z=2
959.02 z=2
1060.56 z=2
366.14 z=1
1754.98 z=1
1917.03 z=1
o
[-CH6O3]⁺ 138.06
[-C2H4O2]⁺ 144.07
[-2H2O]⁺ 168.07
[-C2H4O2]⁺ 126.06
204.09
Ratio intensity = (138.06+168.07)/(126.06+144.07) = >1
b3-H2O 310.21
b3 328.22
366.14
p
1141.58²⁺
ETciD-MS²
I T I L G S N L G V Q A G D I Q R
z15 z14 z13 z12 z11 z10 z9 z8 z7
1141.58 z=2
z9 927.48 z=1
z7 771.39 z=1
z11 1154.61 z=1
z15 1524.83 z=1
1754.98 z=1
1917.03 z=1
z10 1040.56 z=1
z12 1241.64 z=1
z14 1411.74 z=1
z8
z13

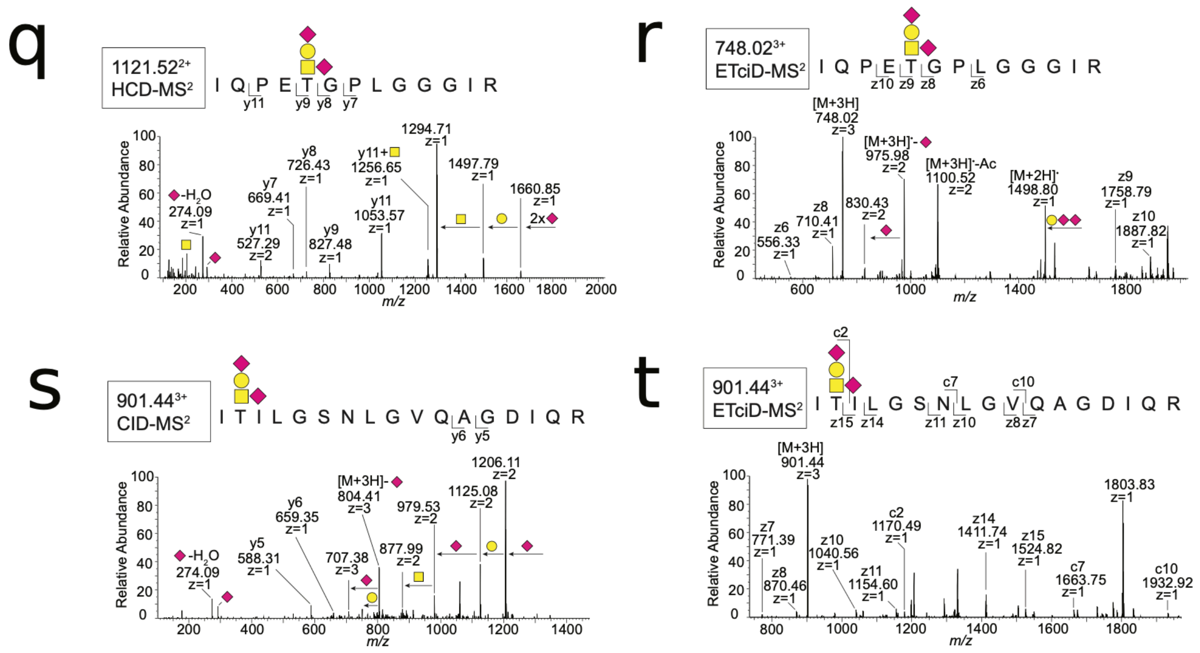

**Fig. S11. Annotated MS/MS spectra of tryptic glycopeptides from sfGFP reporters expressed in glycoengineered HEK293 cell lines.** Precursor ion monoisotopic mass, charge state, and fragmentation mode (HCD, CID or ETciD) are indicated in each panel. Peptide sequences are shown with O-glycan modifications; glycan positions indicated mapped sites (by ETciD-MS2) for each glycopeptide. Glycans are illustrated according to the SFG notation (green circle for mannose, yellow circle for galactose, blue square for N-acetylglucosamine, yellow square for N-acetylgalactosamine, red triangle for fucose and pink diamond for sialic acid [NeuAc]). **a-b,** HCD- and ETciD-MS2 spectra mapping the O-Man modified AGGTC*LTLEGQSLSVGTSR peptide from RON IPT2 (asterix indicates carbamidomethyl modification of cysteine). **c-d,** HCD- and ETciD-MS2 spectra mapping the O-Man modified ITILGSNLGVQAGDIQR peptide from PLXNB2 IPT1. **e-h,** CID- and ETciD-MS2 spectra mapping sialylated and fucosylated core-M1 glycoforms of ITILGSNLGVQAGDIQR. **i-l,** HCD-, CID- and ETciD-MS2 spectra mapping core-M3 glycoforms of ITILGSNLGVQAGDIQR. Panel **j** shows the B-type fragment at m/z 407.17 (HexNAc-HexNAc), and panel **k** shows the oxonium-ion profile, intensities and ratio used to assign the GalNAc epimer, as previously described [13]. **m-p,** HCD-, CID- and ETciD-MS2 spectra mapping additional core-M1 glycoforms, oxonium-ion profile, intensities and ratio used to assign the GalNAc epimer, as previously described [13]. **q-r,** HCD- and ETciD-MS2 mapping the NeuAc-Gal(NeuAc)GalNAc-modified IQPETGPLGGGIR sequence from PLXNB2 IPT1. **s-t,** HCD- and ETciD-MS2 spectra mapping the NeuAc-Gal(NeuAc)GalNAc-modified ITILGSNLGVQAGDIQR peptide from PLXNB2 IPT1.

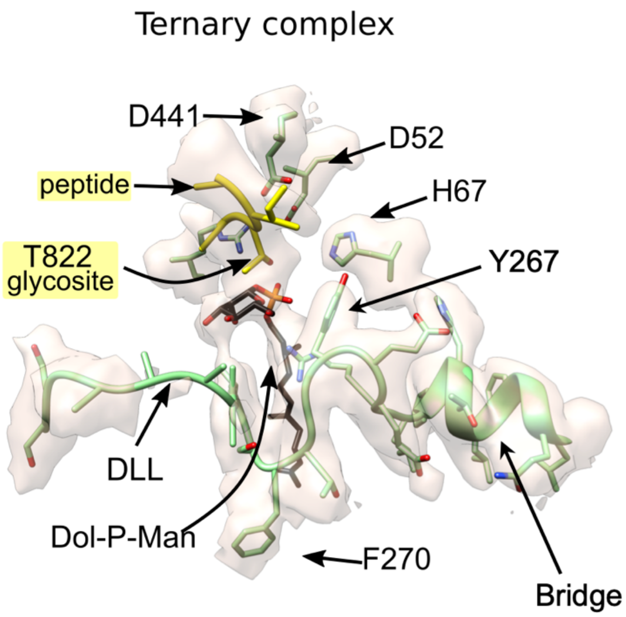

**Fig. S12. Active site architecture in the TMEM260 ternary complex.** Close-up of the active site of TMEM260 bound to the native donor Dol-P-Man and the PLXNB2 IPT1 acceptor peptide (3.1 Å resolution), shown in the same orientation as in Fig. S6. In the ternary complex, D441 is positioned adjacent to the PLXNB2 IPT1 glycosylation site, consistent with a catalytic role, whereas D52 is displaced from the sugar relative to its donor-engaged configuration in the binary complexes (compare with Fig. S6). The donor-loading loop (DLL) is comparatively less well resolved, consistent with increased flexibility in the acceptor-bound state.

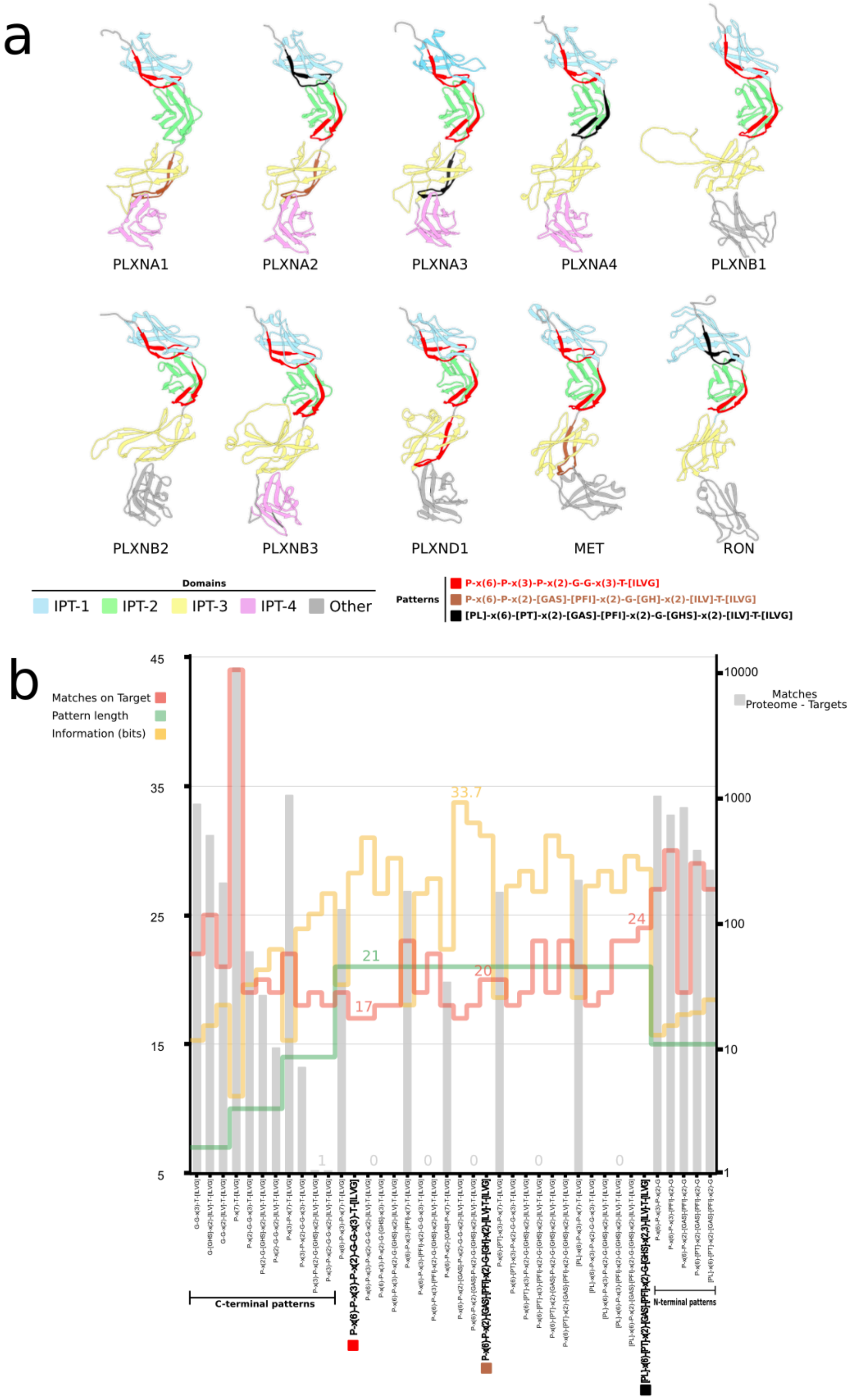

**Fig. S13. Analysis of candidate TMEM260 sequons. a,** AlphaFold-predicted structures of reported human TMEM260 substrates (UniProt) showing extracellular stalk regions containing IPT domains. Candidate sequences are highlighted according to PROSITE pattern matches using cumulative color coding (red ⊂ brown ⊂ black). **b,** Performance of PROSITE-derived sequon patterns. Red indicates matches within the reported substrates in **a**; green, indicates pattern length; yellow, indicates Shannon information content (left axis). Grey bars indicate proteome-wide matches in excess of those in the reported targets (right axis). Selected patterns are labelled. Patterns enriched for three interspaced prolines flanking the A and A′ strands and the A′–B loop GG motif characterize the C-terminal classes (left). N-terminal patterns lacking the glycosite-containing A/A′/partial B strands are shown on the right. Intermediate 21-residue patterns corresponding to the A-B peptide region observed in the ternary complex are shown. Overall, many patterns generate numerous extraneous proteome-wide matches, particularly shorter ones, indicating that 21-residue motifs are necessary but not sufficient for target discrimination. Loss of key sequence features (e.g., the GG motif) also increases false positives, consistent with a minimum information threshold for specificity across both the glycosite/Rl interface and the acceptor-bound TPR region. Given that a defined 21-residue peptide carries ~91 bits of information, the cumulative Shannon information suggests that ~28 bits are required to recover known TMEM260 targets accurately. Together, these analyses support recognition of an interspersed sequon, broad enough to accommodate diverse targets, yet sufficiently constrained to exclude most other proteins.

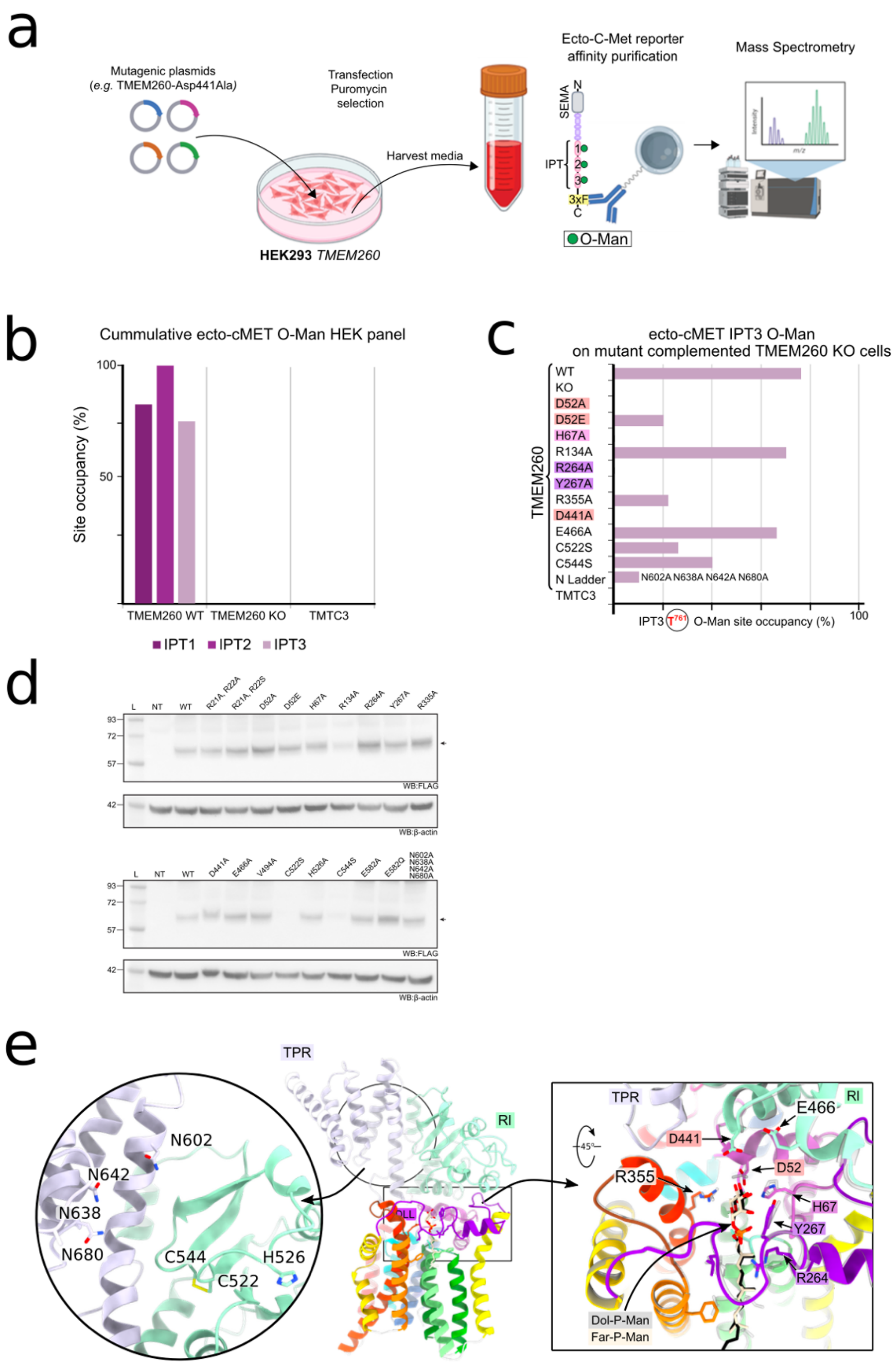

**Fig. S14. O-mannosylation activity of TMEM260 variants. a,** TMEM260-knockout (KO) HEK293 cells expressing a soluble FLAG-tagged ecto-cMET reporter were transfected with wild-type (WT) or mutant TMEM260 constructs, or with TMTC3 as a specificity control, and selected with puromycin. The secreted reporter was affinity-purified using anti-FLAG resin and analyzed by bottom-up mass spectrometry. **b,** All three IPT domains of the ecto-cMET reporter are modified with O-Mannose, with IPT3 selected for quantitative analysis. O-mannosylation is abolished in TMEM260-KO cells and restored upon re-expression of WT TMEM260, in agreement with previous glycoproteomic studies. **c,** Site-resolved glycoproteomic analysis reveals a pronounced loss of activity for variants affecting the catalytic region and the asparagine ladder within the ER-luminal domain. **d,** Immunoblot analysis of TMEM260 variants shows robust expression for most constructs, with a subset exhibiting reduced stability that correlates with diminished O-mannosylation detected by mass spectrometry. **e,** Engineered mutations mapped onto the TMEM260:Far-P-Man binary complex, highlighting their spatial relationship to the donor-binding pocket and acceptor-recognition regions.

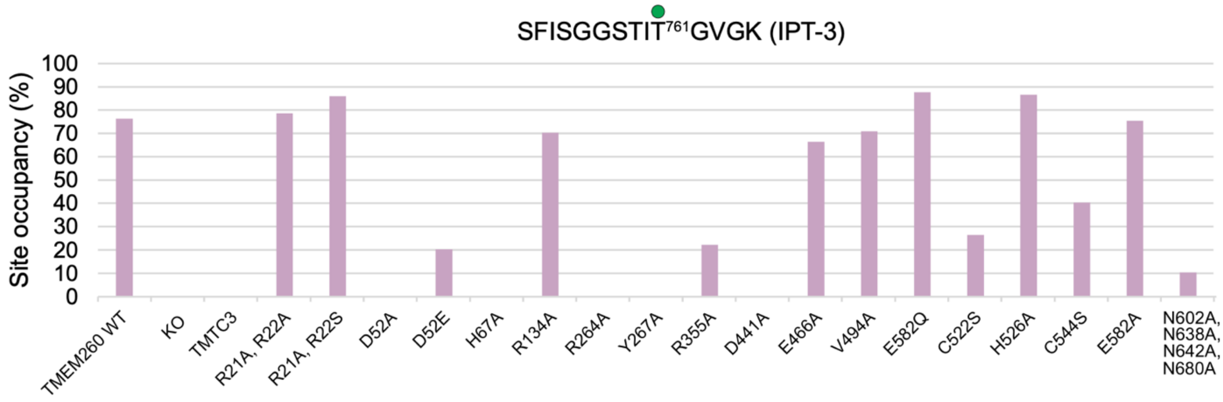

**Fig. S15. O-mannosylation activity of TMEM260 variants in HEK293 cells.** The O-mannosylation activity of TMEM260 variants in HEK393 cells was assayed by glycoproteomic analysis of a FLAG-tagged ecto-cMET reporter substrate carrying a T761 glycosylation site.

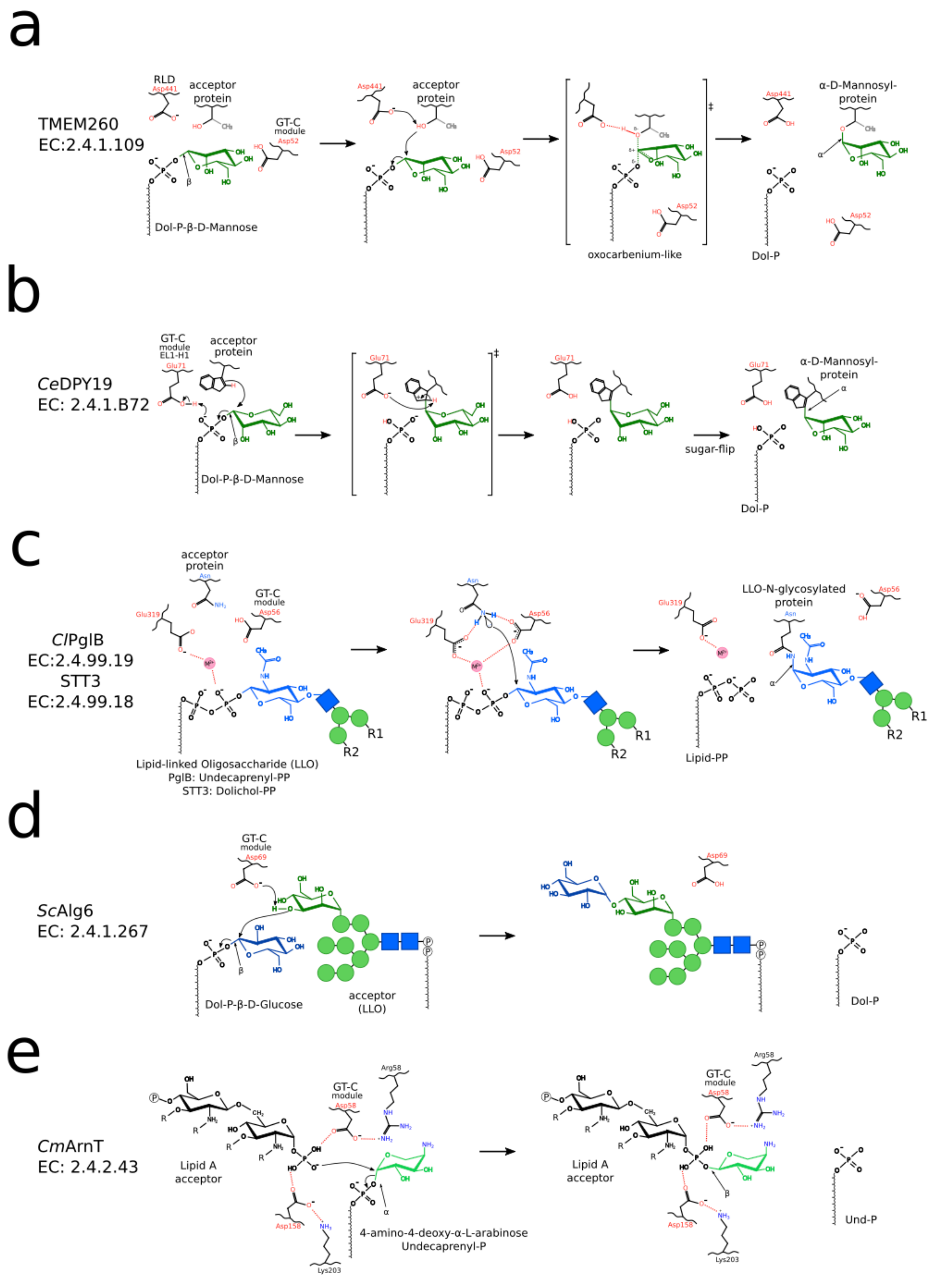

**Fig. S16. Proposed catalytic mechanism of TMEM260 compared with representative GT-C enzymes. a,** Proposed SN2-like mechanism for TMEM260. In the ternary complex, D441 from the Rl domain is positioned adjacent to the acceptor glycosite and is proposed to act as the catalytic base, while D52 from the GT-C module contributes to donor positioning and orientation.

Deprotonation of the threonine hydroxyl enables nucleophilic attack on the C1 of dolichol-β-D-mannosyl phosphate, yielding an α-O-mannose product with anomeric inversion. **b,** Proposed mechanism of the C-mannosyltransferase CeDPY19, involving electrophilic aromatic substitution at the Trp C2 position and inversion at the donor mannose anomeric center. **c,** OST-mediated N-glycosylation, in which a 'twisted amide' activates the asparagine side chain for nucleophilic attack on the anomeric carbon of GlcNAc, forming a β1-N linkage with inversion. **d,** Alg6 mechanism, where D69 deprotonates the C3 hydroxyl of the terminal mannose on the lipid-linked oligosaccharide, promoting nucleophilic attack on Dol-P-Glc and transfer with inversion. **e,** ArnT mechanism, in which an active-site Asp positions the lipid A acceptor for an SN2-like attack on the anomeric carbon of L-Ara4N, resulting in stereochemical inversion.

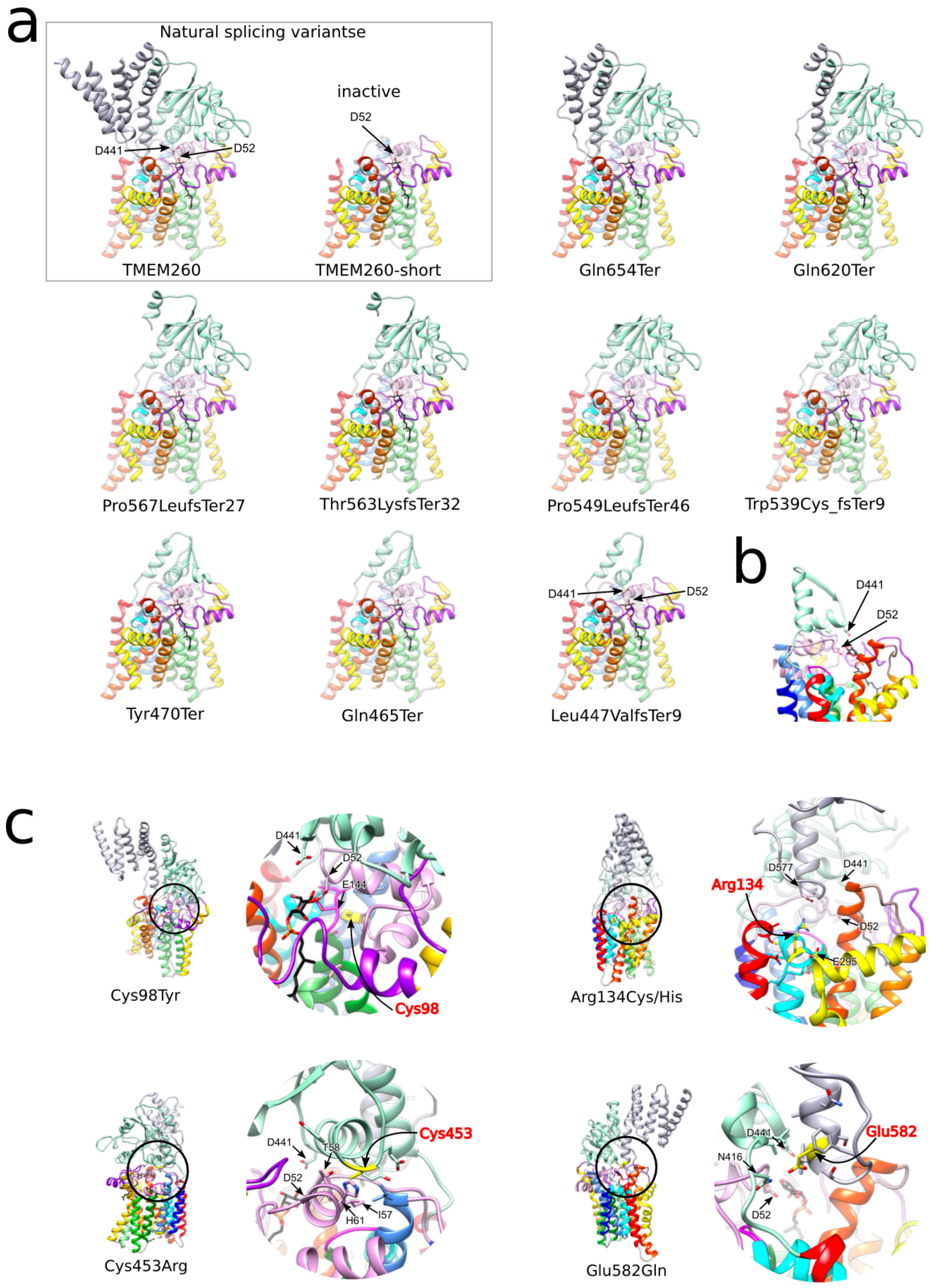

**Fig. S17. Putative structural effects of patient-associated TMEM260 variants linked to congenital heart disease. a,** Reported TMEM260 splice variants and truncations mapped onto the domain architecture. Catalytic residues D52 and D441 are indicated. **b,** Close-up view of the Leu447fs*9 truncation, illustrating the minimal structural framework remaining to support D441 positioning. **c,** Structural mapping of reported missense variants in overall and local views. Most variants lie outside the active site and acceptor-binding cleft, consistent with predominantly allosteric effects. See Supplementary Table 3 for patient details.
